# A comparative analysis of quantitative metrics of root architectural phenotypes

**DOI:** 10.1101/2020.12.01.406827

**Authors:** Harini Rangarajan, Jonathan P. Lynch

## Abstract

High throughput phenotyping is important to bridge the gap between genotype and phenotype. The methods used to describe the phenotype therefore should be robust to measurement errors, relatively stable over time, and most importantly, provide a reliable estimate of elementary phenotypic components. In this study, we use functional-structural modeling to evaluate quantitative phenotypic metrics used to describe root architecture to determine how they fit these criteria. Our results show that phenes such as root number, root diameter, lateral root branching density are stable, reliable measures and are not affected by imaging method or plane. Metrics aggregating multiple phenes such as *total length, total volume, convexhull volume, bushiness index* etc. estimate different subsets of the constituent phenes, they however do not provide any information regarding the underlying phene states. Estimates of phene aggregates are not unique representations of underlying constituent phenes: multiple phenotypes having phenes in different states could have similar aggregate metrics. Root growth angle is an important phene which is susceptible to measurement errors when 2D projection methods are used. Metrics that aggregate phenes which are complex functions of root growth angle and other phenes are also subject to measurement errors when 2D projection methods are used. These results support the hypothesis that estimates of phenes are more useful than metrics aggregating multiple phenes for phenotyping root architecture. We propose that these concepts are broadly applicable in phenotyping and phenomics.

## Introduction

Crop production needs to double by 2050 to provide for the increasing global population (Tilman et al., 2011; Ray et al., 2013; Wise, 2013; FAO, 2017). A major challenge is the identification of efficient crops that cope with climate change and reduce the need for fertilizer and water inputs to make agriculture environmentally sustainable. Root architecture influences water and nutrient uptake, so, selecting and developing efficient crops based on their root system architecture (RSA) has been proposed as a strategy towards a “second green revolution” (Lynch, 2007; Herder et al., 2010; Villordon et al., 2014; Lynch, 2019).

Development of powerful tools in genomic research has resulted in a deluge of genomic information. However, this genomic information cannot be fully exploited for crop improvement unless it is linked to the phenome (Lynch and Brown, 2012; Cobb et al., 2013; Tardieu et al., 2017). In the context of roots, the root phenome is the set of phenes manifested by roots of a plant, where phenes are elementary units of the phenotype; phenes are related to phenotypes as genes are to genotypes (Lynch and Brown, 2012; York et al., 2013). Phenotyping is a bottleneck for breeding and genetic analysis because it is species-specific, labor intensive and environmentally sensitive, unlike genotyping, which is uniform across organisms, highly automated, and increasingly inexpensive (Furbank and Tester, 2011; Lynch and Brown, 2012; Cobb et al., 2013; Atkinson et al., 2019). Phenotyping is especially challenging for roots because of their complexity, plasticity, and inaccessibility. Significant advances are being made in phenotyping methods and technology in an attempt to develop high-throughput platforms. In order to develop efficient strategies to explore the phenome, it is important to clarify what constitutes a phenotype, delineate the key components that comprise a phenotype, and determine the level of resolution at which phenotypic data must be collected. Although an essentially infinite number of measurements may be collected to describe each phenotype, a smaller number of more basic variables may explain most of the important phenotypic variation among genotypes. These basic variables, or *phenes* are the elementary units of the root phenotype and cannot be decomposed to more phenes at the same scale of organization (Lynch and Brown, 2012). Based on this definition, number of axial roots, lateral root branching density (LRBD), root growth angle, root diameter length of different root classes of the root system can be considered as phenes.

Current methods for developing high-throughput phenotyping platforms and identification of relevant quantitative trait loci (QTL) associated with traits of interest are largely based on non-elementary phenotypic metrics. Non-phenes, referred to as phene aggregates in this paper, are aggregate components of the root phenotype and describe the distribution of roots, shape of roots and/or size of the root system. Phene aggregates include several conventionally measured traits including *total root length, total area, total volume*, as well as novel phenotypic metrics such as *convex hull volume, convex hull area, ellipse major axis, ellipse minor axis, ellipse aspect ratio, volume distribution, solidity, bushiness* (Iyer-Pascuzzi et al., 2010; Clark et al., 2011; Cobb et al., 2013; Topp et al., 2013) and metrics which measure the geometry and complexity of root systems such as *fractal dimension* (FD), *fractal abundance* (FA), and *lacunarity* (Fitter and Stickland, 1992; Nielsen et al., 1999; Walk et al., 2004). Aggregate phenotypic metrics (referred to as aggregate metrics) are comprised of phenes, some of these can be measured as a simple aggregate of phenes (e.g. *total length*), some are represented as a function of other aggregates (e.g. *bushiness index, solidity, volume distribution*), some measure shapes resulting from interaction of the constituent phenes (e.g. *Convex hull volume*), and some metrics are complex metrics which measure emergent properties of root architecture and cannot be described as a simple aggregate, shape aggregate or a function of other aggregates (*e.g. Fractal Dimension*).

Estimates of phene aggregates change over time and are phenotype specific. Some phene aggregates increase over time, some remain relatively static and some decrease in value over time (Iyer-Pascuzzi et al., 2010; Zurek et al., 2015). The magnitude of change in estimates of phene aggregates with time also vary greatly. This is because some of the phene aggregates are one-dimensional measurements while some measurements are a function of more than one dimension (Mairhofer et al., 2013). Many phene aggregates are estimates generated from the average values of the 2D projections in a rotational image series (Topp et al., 2013) and are thought to represent 3D root shape accurately. However, which traits can be measured accurately using estimates derived from 2D data and which require 3D representations is poorly understood. Depending on the phenotype, metrics derived from rotated 2D projections of the same 3D root system can vary significantly. This leads to a related question of how much should an aggregate phenotypic metric differ for two phenotypes to be considered distinctly different. Fractal analysis of corn roots have shown that the *FD* of two genotypes can be same but vary in *FA* (Eghball et al., 1993). Root systems with similar *FD* may vary functionally and genotypes can be distinguished when fractal analysis involves *FD, FA* and *lacunarity* (Walk et al., 2004). Aggregate phene metrics estimate the aggregate of multiple phenes. For example, greater rooting depth is an important trait for capture of subsoil N in maize. Greater rooting depth results from a combination of deeper axial root growth angle (Manschadi et al., 2006; Trachsel et al., 2013; Uga et al., 2013), root elongation rate (Manschadi et al., 2008), expression of fewer crown roots (Saengwilai et al., 2014b; Gao and Lynch, 2016), reduced lateral branching density (Postma et al., 2014; Zhan et al., 2015), formation of root cortical aerenchyma (RCA) (Postma and Lynch, 2011; Saengwilai et al., 2014a), reduced cortical file number and increased cortical size (Jaramillo et al., 2013; Chimungu et al., 2014). Each of these phenes are under distinct genetic control and have important interactions with each other. Selection for combination of specific phenes will therefore be much simpler and precise than would selection for root depth itself (Lynch, 2019). Phenes are under more simple genetic control and permit more precise control over the root system architecture (RSA) and so, are more useful for selection for crop breeding (Lynch and Brown, 2012; Lynch, 2019).

In this study, we use the functional-structural plant model *SimRoot* to identify phenotyping metrics that are

- sensitive enough to provide information on the constituent root phenes and their states,
- stable over time and are independent of the time of phenotyping,
- robust to the imaging method *i.e.*, do not vary when measured in the intact 3D root system or when estimated using 2D rotational image series.

Our analysis shows that

- Phene aggregates can be explained by phenes. Different phene aggregates capture different combinations of subtending phenes. However, these metrics do not provide any information or measure of the phene state of the constituent phenes.
- Several combinations of phenes in different states can produce phenotypes which have comparable estimates of phene aggregates. Estimates of phene aggregates are not unique representations of the state of the underlying phenes.
- As the number of phenes captured by an aggregate phenotypic metric increases, the stability of that metric becomes less stable over time.

## 2. Materials and Methods

### 2.1 Simulation of phenotypes

The functional-structural plant model *SimRoot* (Lynch et al., 1997) was used to simulate bean (*Phaseolus vulgaris*) and maize (*Zea mays*) root phenotypes. In *SimRoot*, simulated root system comprises of roots of distinct classes as specified by their root diameters, lateral root branching density, root growth rate and root growth angle in the input parameters. The root growth angle over time depends on the gravitropism. Stochasticity is included in all parameters. The roots are simulated as small connected root segments over time. Co-ordinates corresponding to the root being simulated as well as the root length, volume, area parameters are stored for the simulated root segments as the root grows at specified time points. The root length, area, volume of the root system is estimated by integrating the respective parameters over all root segments. The root image co-ordinates are used to visualize the simulated root system.

The number of roots of different root classes, angle, diameter, lateral root branching density (LRBD) were varied to produce 1500 maize root phenotypes and 1500 bean root phenotypes. The range of values used for each of the root parameter used are given in Supplementary Material 2.

The data corresponding to the simulated root phenotypes were saved during the simulation runs. These data files contained the X, Y, Z co-ordinates of the simulated root system images used to simulate the root as well as data of root length, area, volume etc. of the simulated root segments with their corresponding root class. Roots were allowed to grow without any boundaries so that the growing roots did not touch any boundary surface and so no artifacts were introduced due to mirroring roots. Stochasticity was included in all the simulated parameters. Root growth angle was influenced by root gravitropism. In order to obtain accurate estimates of all the phenotypic traits, elementary and aggregate phenotypic were extracted/calculated from the data of the simulated images.

### 2.2 Measurement of phene and aggregate phene metrics

Estimates of phene metrics were measured from the simulated images. Aggregate phenotypic trait metrics were calculated for intact 3D root systems as well as projections of the roots systems on a 2D plane. The root system was rotated by 20 degrees and the projections on a 2D plane were obtained (Figure 1 and Supplementary Figure S1). The average of the estimates of each metric in all the projected images for each phenotype was used in studies considering 2D projections. The average value was used also in 3D studies where 3D estimates were not obtained including ellipse major axis, ellipse minor axis and ellipse aspect ratio. The phene aggregates estimated and considered in this study, the definitions of these traits and the method of obtaining those metrics from *SimRoot* output is given in Table 1.In order to evaluate how phene metrics and phenotypic trait metrics change over time, root images were obtained every 5 days starting 10 days after germination and metrics obtained for these root systems. This way phenotyping metrics were obtained for 3D root systems, 2D projections of the root systems, and root system images after different periods of growth.

**Table 1:** Aggregate phene metrics, definition and method of obtaining them from *SimRoot* output

| Parameter | 3D | 2D | Description | Measurement |
| --- | --- | --- | --- | --- |
| Total Length | Y | Y | Summed length along the whole root system | Calculated from *SimRoot* output |
| Total Area | Y | Y | Summed surface area of the whole root system | Calculated from *SimRoot* output |
| Total Volume | Y | Y | Summed volume of the whole root system | Calculated from *SimRoot* output |
| Maximum width | Y | Y | Maximum horizontal width of the whole root system | Calculated using minimum enclosing circle algorithm in R |
| Maximum depth | Y | Y | Maximum vertical depth of the whole root system | Calculated from *SimRoot* output |
| Median no. of roots | Y | Y | Median no. of roots from root counts | Calculated from *SimRoot* output |
| Maximum no. of roots | Y | Y | No. of roots at the 84th percentile of a sorted list (smallest to largest) of root counts | Calculated from *SimRoot* output |
| Bushiness | Y | Y | Ratio of the maximum no. of roots to the median no. of roots | Calculated from *SimRoot* output |
| Volume distribution | Y | Y | Ratio of the volume of the root system contained above one-third depth of the root system to the volume of the root system contained below one-third depth of the root system | Calculated from *SimRoot* output |
| Convex hull volume | Y | Y | Volume of the convex hull that encompasses the whole root system | Obtained using `ConvHulln` function in R |
| Convex hull area | Y | Y | Surface Area of the convex hull that encompasses the whole root system | Obtained using `ConvHulln` function in R |
| Solidity | Y | Y | Ratio of volume to convex hull volume | Calculated |
| Major Ellipse axes | Y | N | Length of major axis of an ellipse best fit to overall shape and size of root system | Obtained using minimum volume enclosing ellipse algorithm in R |
| Minor Ellipse Axes | Y | N | Length of minor axis of an ellipse best fit to overall shape and size of root system | Obtained using minimum volume enclosing ellipse algorithm in R |
| Ellipse axis aspect ratio | Y | N | Ratio of major axis of ellipse to minor axis | Calculated from minor ellipse axes and major ellipse axes |
| Fractal Dimension (FD) | Y | Y | Measure of root complexity. Fractal dimension expresses the space filling properties of a structure (e.g. root system) and is associated with branching pattern | Obtained using box count code written in R |
| Fractal Abundance (FA) | Y | Y | Measure of root complexity. Fractal abundance is associated with the volume of space explored | Obtained using box count code written in R |

**Figure 1:**
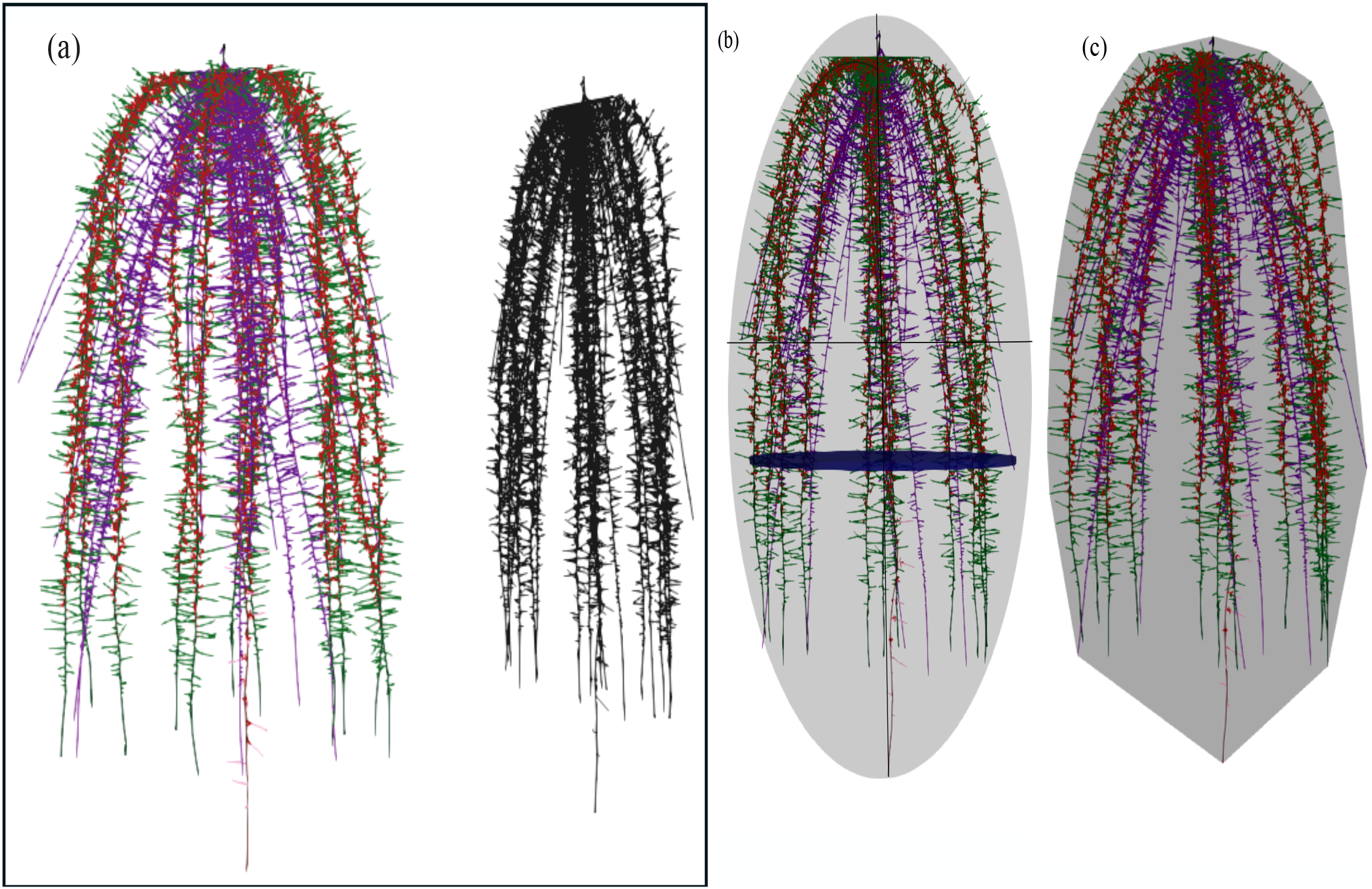
Representation of 2D projection of 3D root system (a) Visualization of maximum width, ellipse major and minor axis (b) Convex hull volume of a 3D root system.

### 2.3 Random forest analysis

Data obtained from 3D root systems were analyzed using Random Forest regression. For metrics where 3D metric data were not available (ellipse minor axis, ellipse major axis and ellipse aspect ratio), the average value of the aggregate phenotypic trait from 2D rotational series was used. Random Forest, is a nonparametric technique derived from classification and regression trees (CART). Random Forest consists of a combination of many trees, where each tree is generated by boot-strap samples, leaving about a third of the overall sample for validation (the out-of-bag predictions – OOB). Each split of the tree is determined using a randomized subset of the predictors at each node. The final outcome is the average of the results of all the trees (Breiman, 2001; Cutler et al., 2007). It uses the OOB samples (independent observations from those used to grow the tree) to calculate error rates and variable importance, no test data or cross-validation is required. However, this method does not calculate regression coefficients nor confidence intervals (Cutler et al., 2007). It allows the computation of variable importance measures that can be compared to other regression techniques. The R package Random Forest was employed for the data analyses, with ntree =1000 and mtry =8. Random forest regression was used with each aggregate phenotypic metric as the dependent variable and the input variables as the independent variables to identify the most important variables. The selection of the most relevant variables to include in the final model was done by ranking the variables according to their importance and excluding the least important variables. The variable importance measure, the mean decrease in accuracy (%IncMSE) was used for selecting the important variables. Variable importance is measured by mean squared error of a variable p, which is averaged increase in prediction error among all regression trees when the OOB data for variable p is randomly permuted. If variable p is important there will be an increase in prediction error. Random forest was conducted 50 times and 90 percentile from distribution of mean squared error as the significance threshold of individual variables. The variables thus chosen were used to run a reduced variable model of the original random forest model for each aggregate metric. The reduced variable models were deemed acceptable if the Random Forest trained upon the most important descriptors gave a fit to the data set which was similar or better than that trained upon all variables.

### 2.4 Variation in estimates of phene aggregate metrics

One aspect of the study was to find if estimates of aggregate phenotypes were a unique representation of the phenes. To address this, a representative phenotype was chosen for the maize root system and phenotypes varying by less than 1 % of an aggregate phenotypic trait a shape phenotypic trait (Convex hull Volume) and a complex phenotypic trait (FD) were chosen to find if the phenes constituting the phenotype varied when the aggregate phenotypic trait was similar. In an alternate approach, the estimates of convex hull volume and FD of bean root phenotypes with differences in basal root whorl number and root growth angles with distinct functional value (Rangarajan et al., 2018) were studied.

### 2.5 Estimates of phene and aggregate phene metrics obtained from 2D projections

In order to study the variation in metrics estimated in 2D rotational image series, the coefficient of variation for each phenotype for each phenotypic trait metric was calculated from 2D projections of the root system and the phenotypic metrics were compared.

### 2.6 Estimates of phene and aggregate phene metrics over time

Root system image data were saved every 5 days from day 10 to day 40 of growth and the 3D estimates of the phenes and phene aggregates were collected.

## 3. Results

Different bean and maize phenotypes were simulated by varying input parameters in *SimRoot*.

### 3.1 Variation in simulated phenotypes

The estimates of all phenotypes were min-max scaled and the phenotypes were clustered by hierarchical cluster analysis of the phenotypes based on their phenes. The results of our study are based on a wide array of phenotypes. Phenotypes included in the study had vastly different phenotypes and differed in few or many phenes. The heatmap in Figure 2(a) shows a small subset of data: the relative values of the bean phenes in a few phenotypes (rows) and the corresponding phenotypes in Figure 2(b). Phenotype 1 had very shallow basal root growth angle compared to phenotype 2 while phenotypes 8 and 9 had deep basal root growth angles. Phenotype 7 had more basal roots than the other phenotypes. Phenotypes 5 and 6 differed in the basal root branching density as well as basal root angle. The heatmap in Figure 3(a) shows a small subset of data: the relative values of maize phenes in a few phenotypes (rows) and the corresponding phenotypes in Figure 3(b). Phenotypes 2 and 3 differed in the number of nodal roots with phenotype 2 having more nodal roots than phenotype 3. Phenotypes 4 and 6 had similar primary root lateral branching but phenotype 6 had no seminal roots while phenotypes 4 had 5 seminal roots. Phenotypes 8 and 9 differed in the number of seminal roots as well as seminal root LRBD and the number of nodal roots. The heatmap of all bean root phenotypes and representative phenotypes considered in this study is included in Supplementary Figure S2(a) and S2(b). A similar heatmap for maize root phenotypes are presented in Supplementary Figure S3(a) and S3(b) respectively.

**Figure 2:**
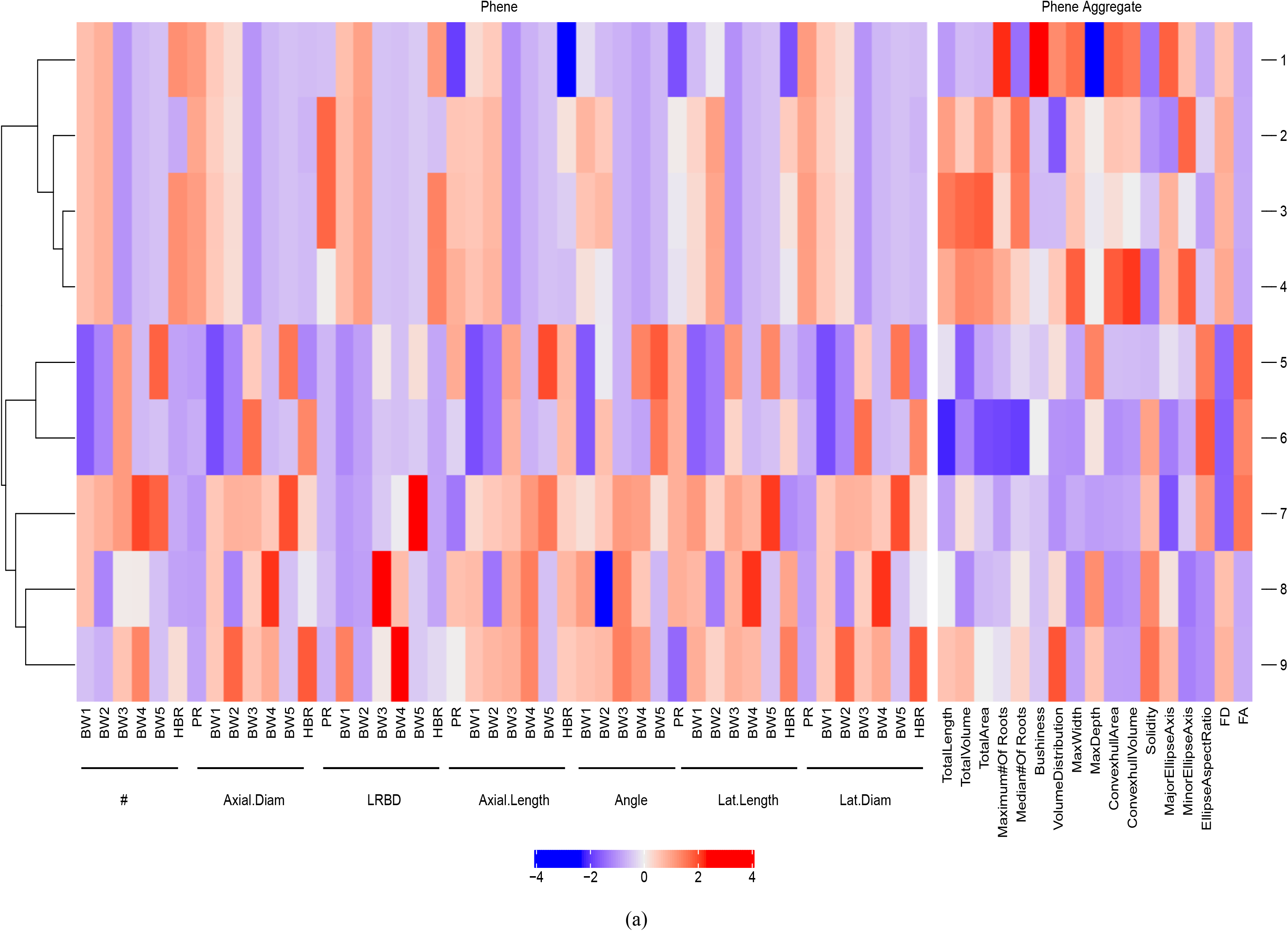
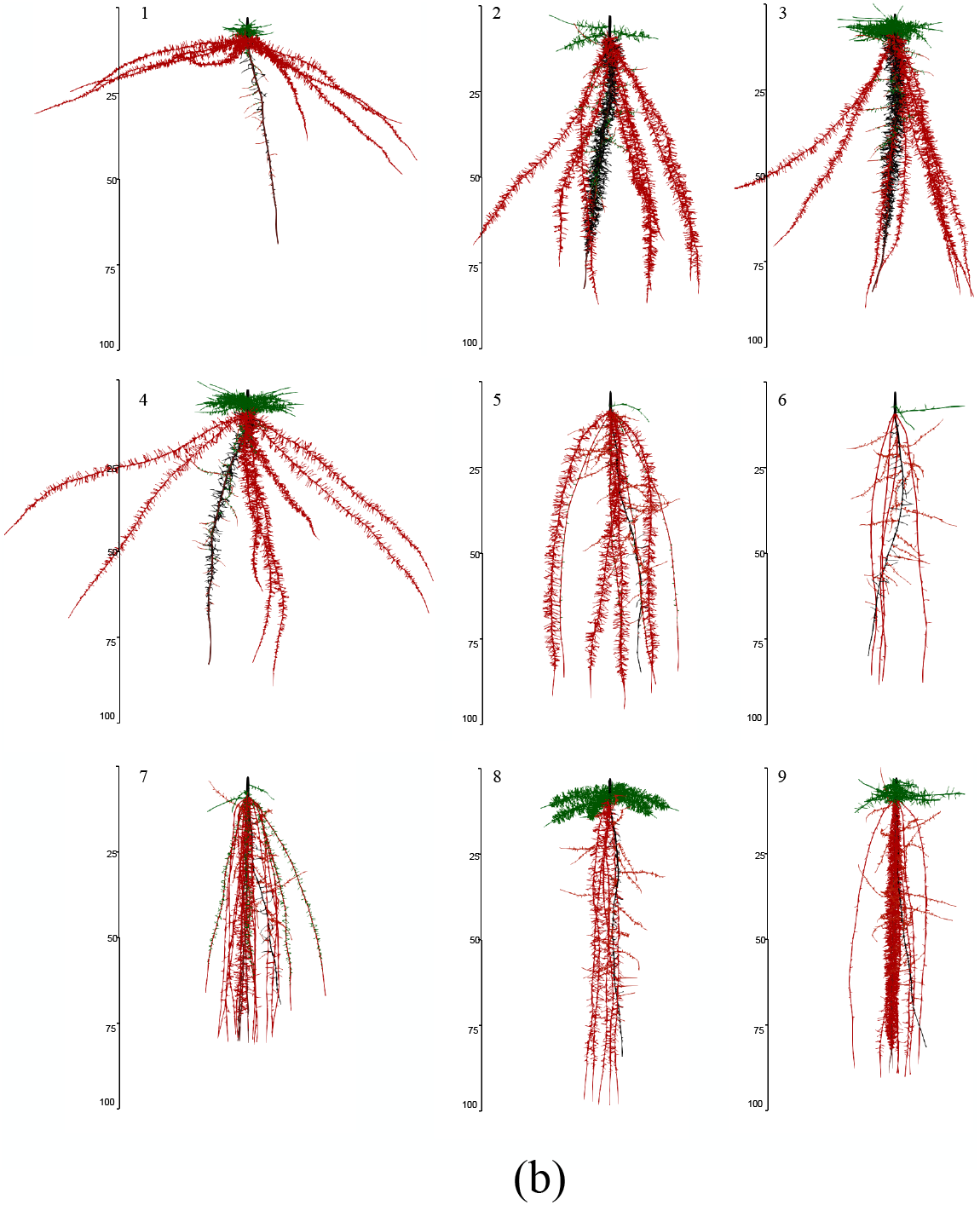
Cluster heatmap of phenotypic traits. Hierarchical clustering of a few phenotypes was generated using Spearman correlation coefficient of max-min scaled phene values of bean phenotypes at 40 days (a). The color scale indicates the magnitude of the trait values (blue, low value; red, high value). The numbers indicated on the heatmap refer to a phenotype in the specific row of the heatmap. The corresponding phenotypes are visualized in (b). Primary roots are in black; basal roots are in red; hypocotyl-borne roots are in green. # - number of axial roots; Axial.Diam - axial root diameter; LRBD - lateral root branching density; Lat.Length - lateral root length; Lat.Diam - lateral root diameter. BW1 - basal roots at whorl 1; BW2 - basal roots at whorl 2; BW3 - basal roots at whorl 3; BW4 - basal roots at whorl 4; BW5 - basal roots at whorl 5; HBR - hypocotyl-borne roots.

**Figure 3:**
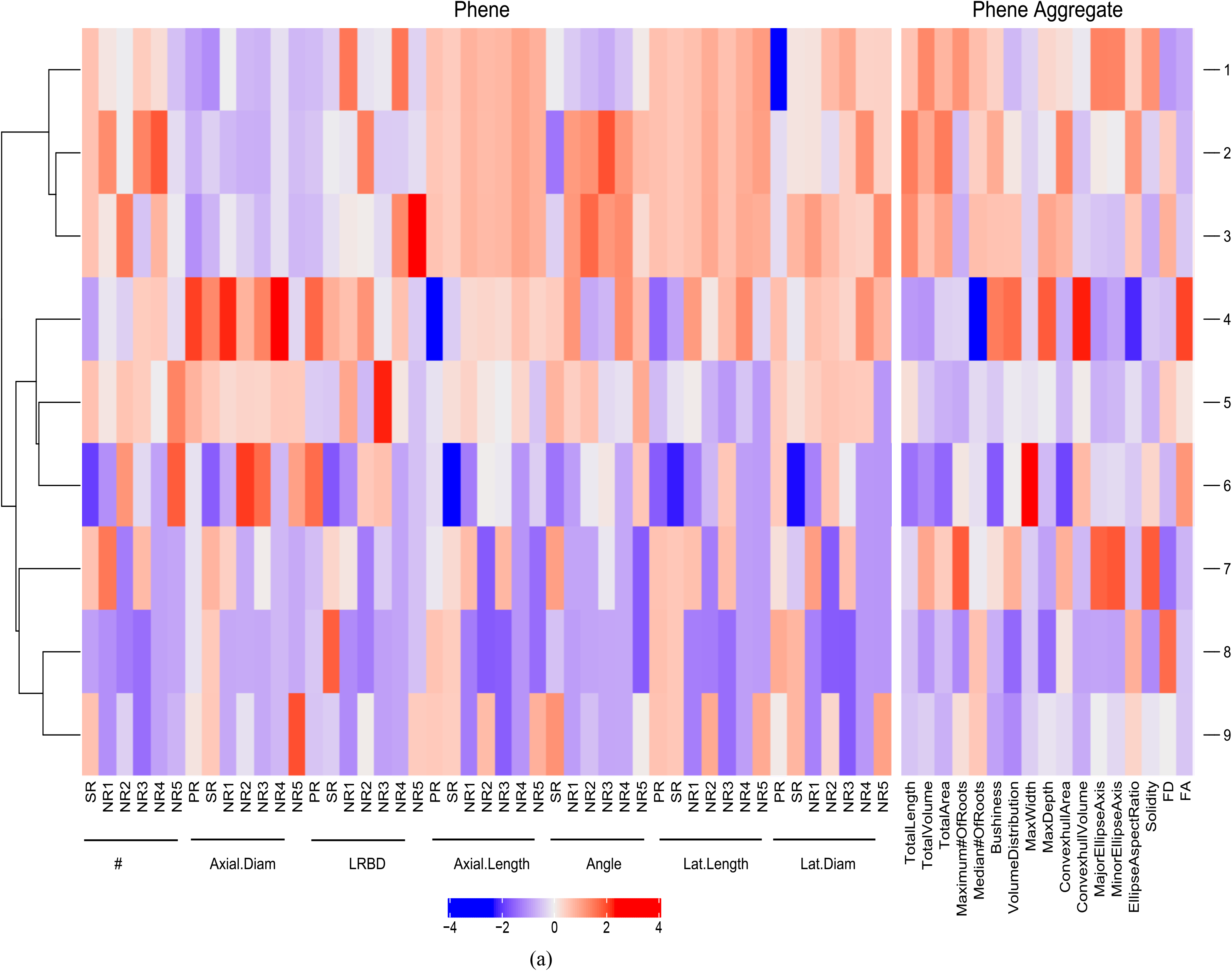
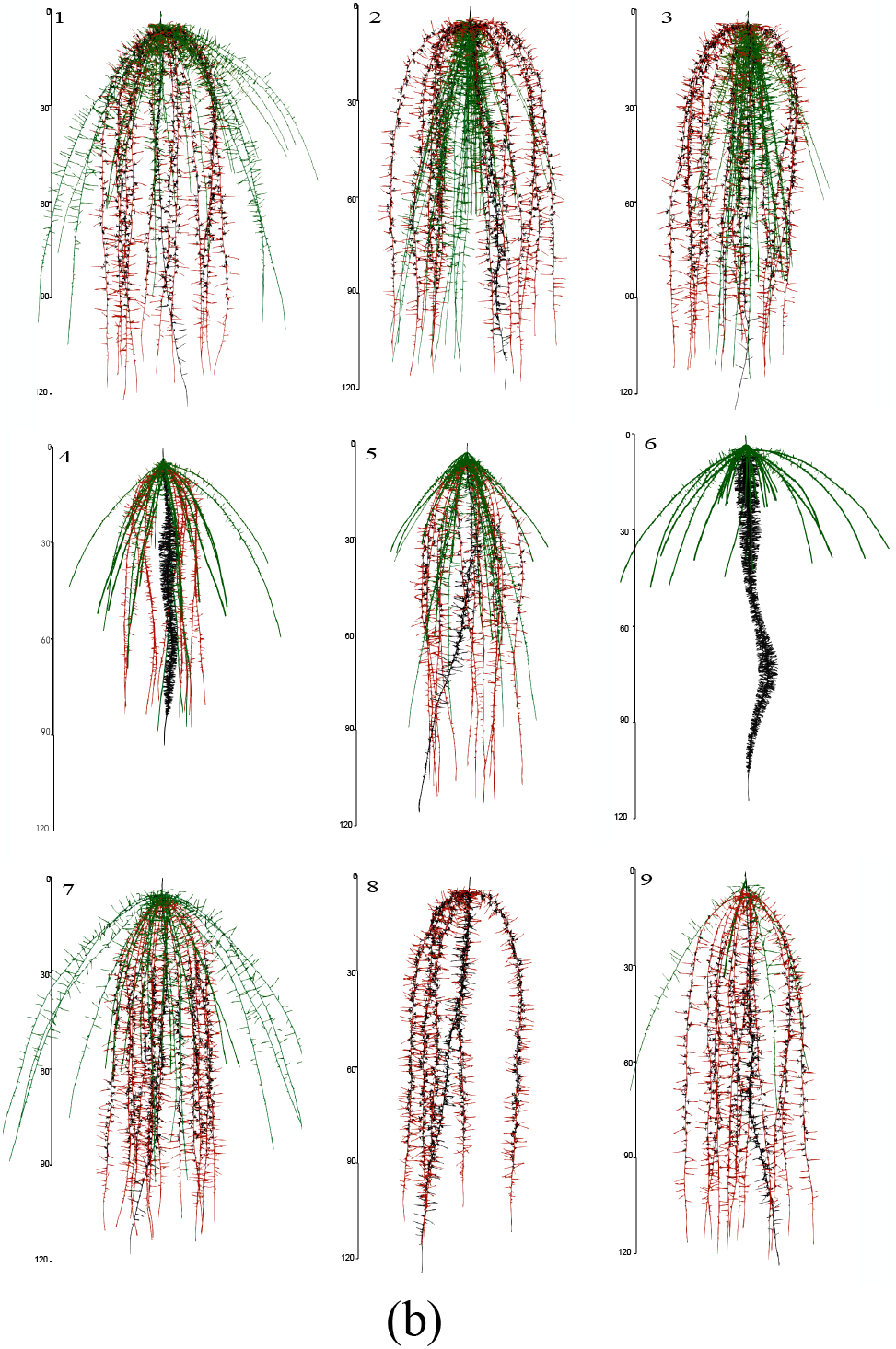
Cluster heatmap of phenotypic traits. Hierarchical clustering of a few phenotypes was generated using Spearman correlation coefficient of max-min scaled phene values of maize phenotypes at 40 days (a). The color scale indicates the magnitude of the trait values (blue, low value; red, high value). The numbers indicated on the heatmap refer to the phenotype in the specific row of the heatmap. The corresponding phenotypes are visualized in (b). Primary roots are in black; seminal roots are in red; nodal roots are in green. # - Number of roots; Axial.Diam - axial root diameter; LRBD - lateral root branching density; Axial.Length - axial root length; Lat.Length-lateral root length; Lat.Diam - lateral root diameter. NR1 - nodal roots at position 1; NR2 - nodal roots at position 2; NR3 - nodal roots at position 3; NR4 - nodal roots at position 4; SR - seminal roots; PR - primary root.

### 3.2 Correlation among phenotypic metrics

Strong correlations were found among the phenes (Figure 4(a) and Figure 4(b)), in the bean root system as well as the maize root system. Axial root length was negatively correlated with diameter, number and LRBD of basal roots in bean and nodal roots in maize root system. The primary axial root length and seminal axial root length was negatively correlated with diameter of the primary root, seminal root axial root length was also negatively correlated with nodal root LRBD. Phenotypes with longer axial roots had greater *maximum width, maximum depth, convex hull area, convex hull volume, major ellipse axis, minor ellipse axis* but smaller values for *solidity* (Figure 4(b). *Solidity* was positively correlated with diameter and number of basal roots in bean. Strong correlations also exist between aggregate phenotypic trait metrics*. Major ellipse axis* positively correlated with *maximum depth*. *Convex hull area*, *convex hull volume*, *minor ellipse axis* and *maximum width* are highly positively correlated with each other. but are negatively correlated with *solidity* (Figure 4).

**Figure 4:**
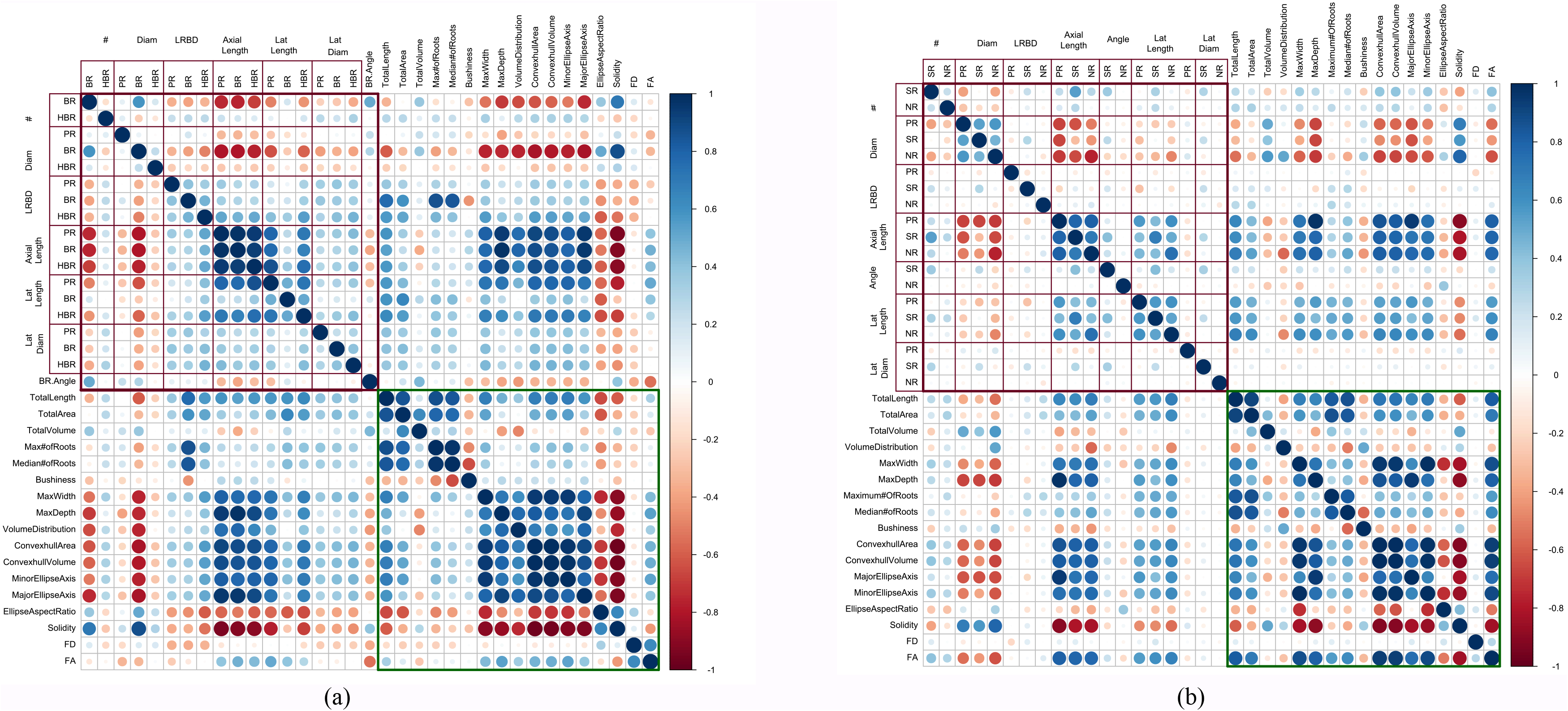
Phenotypic trait relationship. Correlation matrix of phenes and phene aggregates evaluated for bean root phenotypes (a). Correlation matrix of phenes and phene aggregates evaluated for maize root phenotypes (b). The color scale indicates Spearman correlation coefficients between two traits. Color intensity and size of the circle are proportional to the correlation coefficients between the two traits. Correlations between phenes are indicated by the points in the red box, the green box contains the correlations between phene aggregates. BR - basal roots; HBR - hypocotyl-borne roots; PR - primary root; SR - seminal roots; NR - nodal roots; # - number of axial roots; Diam - axial root diameter; LRBD - lateral root branching density; Axial Length - axial root length; Lat Length - lateral root length; Lat Diam - lateral root diameter; FD - fractal dimension; FA - fractal abundance.

### 3.3 Random forest analysis: Different phenes are important in determining the estimate of different aggregate phenes

The results of the random forest analysis are shown in Table 2. Reduced variable models created with Random Forest show proportion of explained variance (R^2^) between 80 % and 99 % for models with all aggregate phenotypic metric except *bushiness index*, which had 62 % in bean and 41% in maize; and *FD* which had R^2^ of 67 % in bean and 20 % in maize. The most important variables for each aggregate phenotype for the bean and maize models are summarized in Table 3. The variables have been summarized based on the phene the variable represents.

**Table 2:**
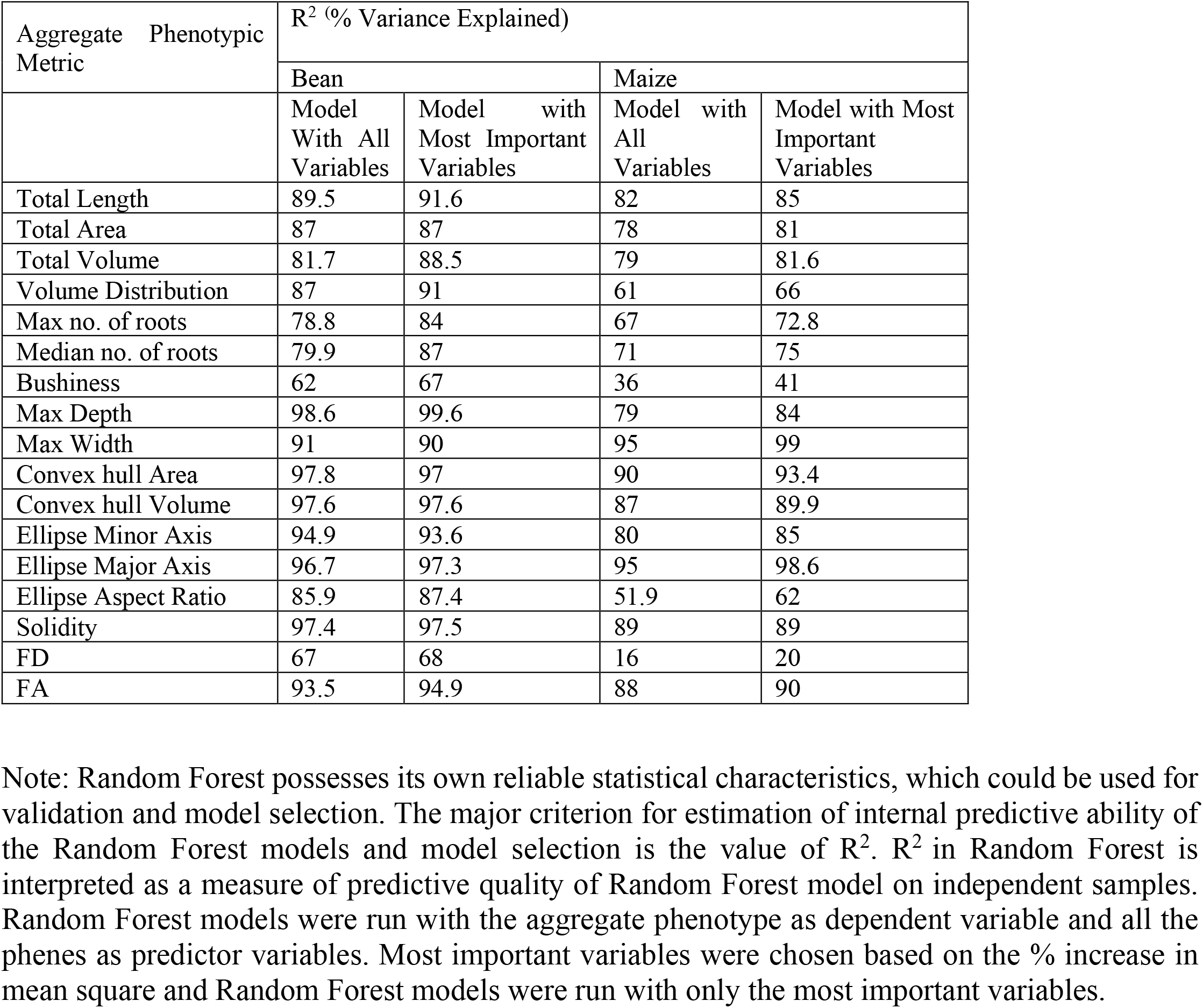
Results of regression models created with random forest. The R^2.^ values of Random Forest model with entire set of variables and those with only most important variables are presented for the bean and maize aggregate phene metrics.

| Aggregate Phenotypic Metric | $R^2$ (% Variance Explained) | | | |
| --- | --- | --- | --- | --- |
|  | Bean |  | Maize |  |
|  | Model With All Variables | Model with Most Important Variables | Model with All Variables | Model with Most Important Variables |
| Total Length | 89.5 | 91.6 | 82 | 85 |
| Total Area | 87 | 87 | 78 | 81 |
| Total Volume | 81.7 | 88.5 | 79 | 81.6 |
| Volume Distribution | 87 | 91 | 61 | 66 |
| Max no. of roots | 78.8 | 84 | 67 | 72.8 |
| Median no. of roots | 79.9 | 87 | 71 | 75 |
| Bushiness | 62 | 67 | 36 | 41 |
| Max Depth | 98.6 | 99.6 | 79 | 84 |
| Max Width | 91 | 90 | 95 | 99 |
| Convex hull Area | 97.8 | 97 | 90 | 93.4 |
| Convex hull Volume | 97.6 | 97.6 | 87 | 89.9 |
| Ellipse Minor Axis | 94.9 | 93.6 | 80 | 85 |
| Ellipse Major Axis | 96.7 | 97.3 | 95 | 98.6 |
| Ellipse Aspect Ratio | 85.9 | 87.4 | 51.9 | 62 |
| Solidity | 97.4 | 97.5 | 89 | 89 |
| FD | 67 | 68 | 16 | 20 |
| FA | 93.5 | 94.9 | 88 | 90 |

Note: Random Forest possesses its own reliable statistical characteristics, which could be used for validation and model selection. The major criterion for estimation of internal predictive ability of the Random Forest models and model selection is the value of $R^2$ . $R^2$ in Random Forest is interpreted as a measure of predictive quality of Random Forest model on independent samples. Random Forest models were run with the aggregate phenotype as dependent variable and all the phenes as predictor variables. Most important variables were chosen based on the % increase in mean square and Random Forest models were run with only the most important variables.

**Table 3:**
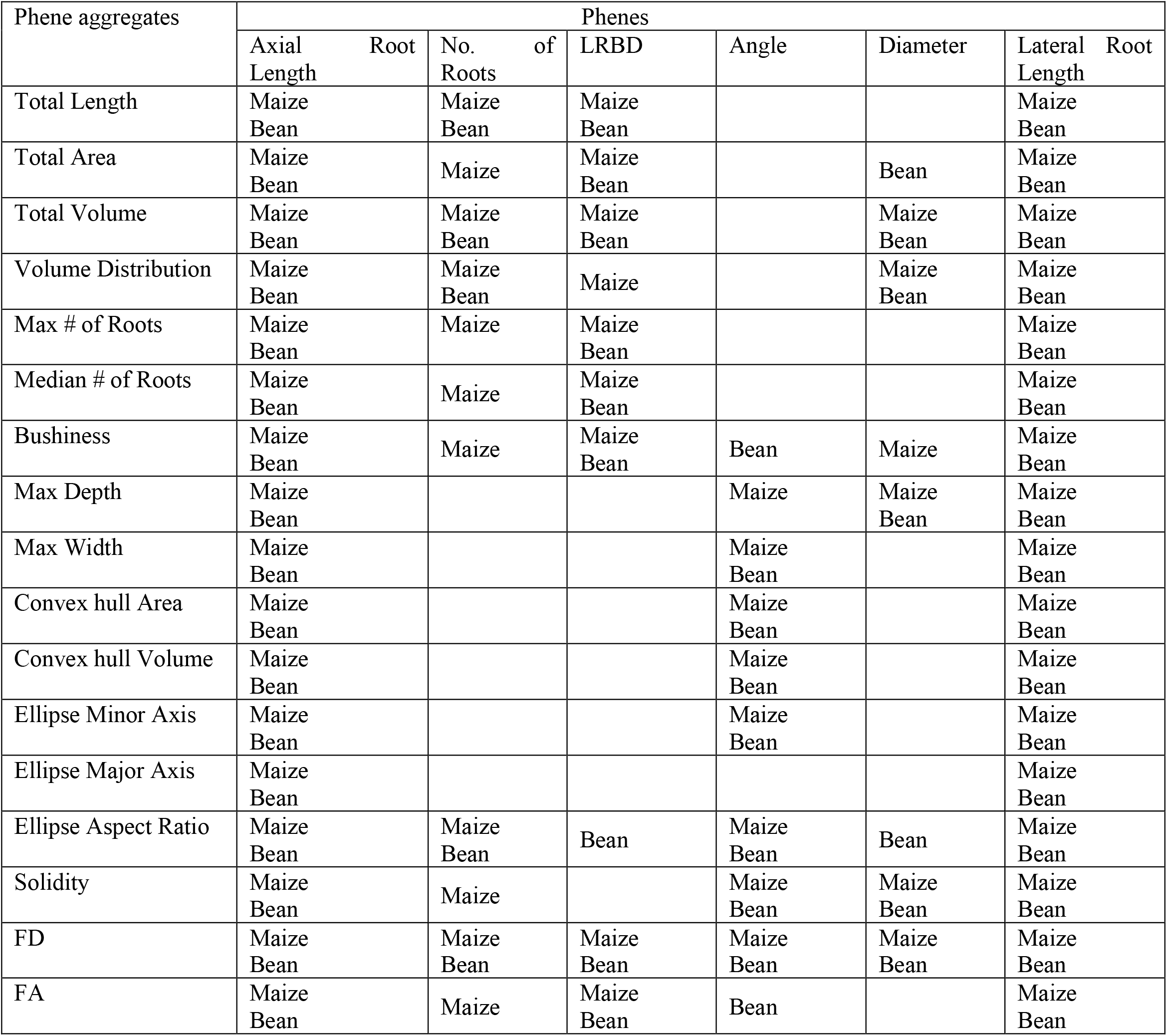
Summary of the most important variables selected by random forest model for each phenotyping metric evaluated for bean root system and maize root system

Among the variables evaluated by the random forest analysis, axial root length and lateral root length were found to be important explanatory variables for all the phene aggregates in both bean as well as maize. Lateral Root Branching Density (LRBD) was found to be an important variable for *total length, total area, total volume, maximum number of roots, median number of roots bushiness index*, *FD* and *FA* in bean as well as maize. LRBD was also important for *volume distribution* in maize root phenotypes and *ellipse aspect ratio* in bean root phenotypes. Number of roots and diameter played important roles in determining the *total area* in maize and bean root systems respectively. Root diameter was an important variable for *total volume, volume distribution, maximum depth, solidity* and *FD* in both bean and maize phenotypes. Diameter was also an important variable in *total area* and *ellipse aspect ratio* in bean and *bushiness index* in maize root phenotypes. Angle was selected as an important variable by the random forest models for *maximum width, convex hull area, convex hull volume, ellipse minor axis, ellipse aspect ratio, solidity* and *FD* for both maize and bean. All the variables evaluated are important for the model with *FD* as the dependent variable.

### 3.4 Estimates of aggregate phene metrics can be similar for phenotypes with different phene states

Even in phenotypes with similar estimates for aggregate phenotypic metrics, the phene states of the constituent phenes varied greatly (Figure 5(a), Figure 5(b)). Phenotypes chosen based on the similarity of aggregate phenotypic metrics had different diameter, LRBD, and number of roots of different classes. Conversely, phenotypes in which phenes exist in different states have similar aggregate phenotypic metrics (Figure 6). Four bean phenotypes that vary only in the number of basal roots and root growth angle were chosen and the estimate of total volume, convex hull volume and FD were compared (Figure 6). Phenotype 1 has one whorl of basal roots with shallow angles, phenotype 2 has one whorl of basal roots with deep angles, phenotype 3 has three whorls with fanned root growth angles. While phenotypes 1 and 2, which vary only in root growth angle, have different estimates for all the three metrics considered (total volume, convexhull volume and FD) phenotypes 1 and 3 have similar estimates for FD (varying by less than 2%) even though they vary in both in number of basal roots as well as root growth angles. Similarly, phenotype 4 has four whorls with fanned angles and differs from phenotype 3 and phenotype 1 in number of basal roots as well as root growth angle, but varies in the estimates of total volume by 1% and 16 % respectively; and in the estimate of convexhull volume by 1% and 4% respectively (Figure 6).

**Figure 5:**
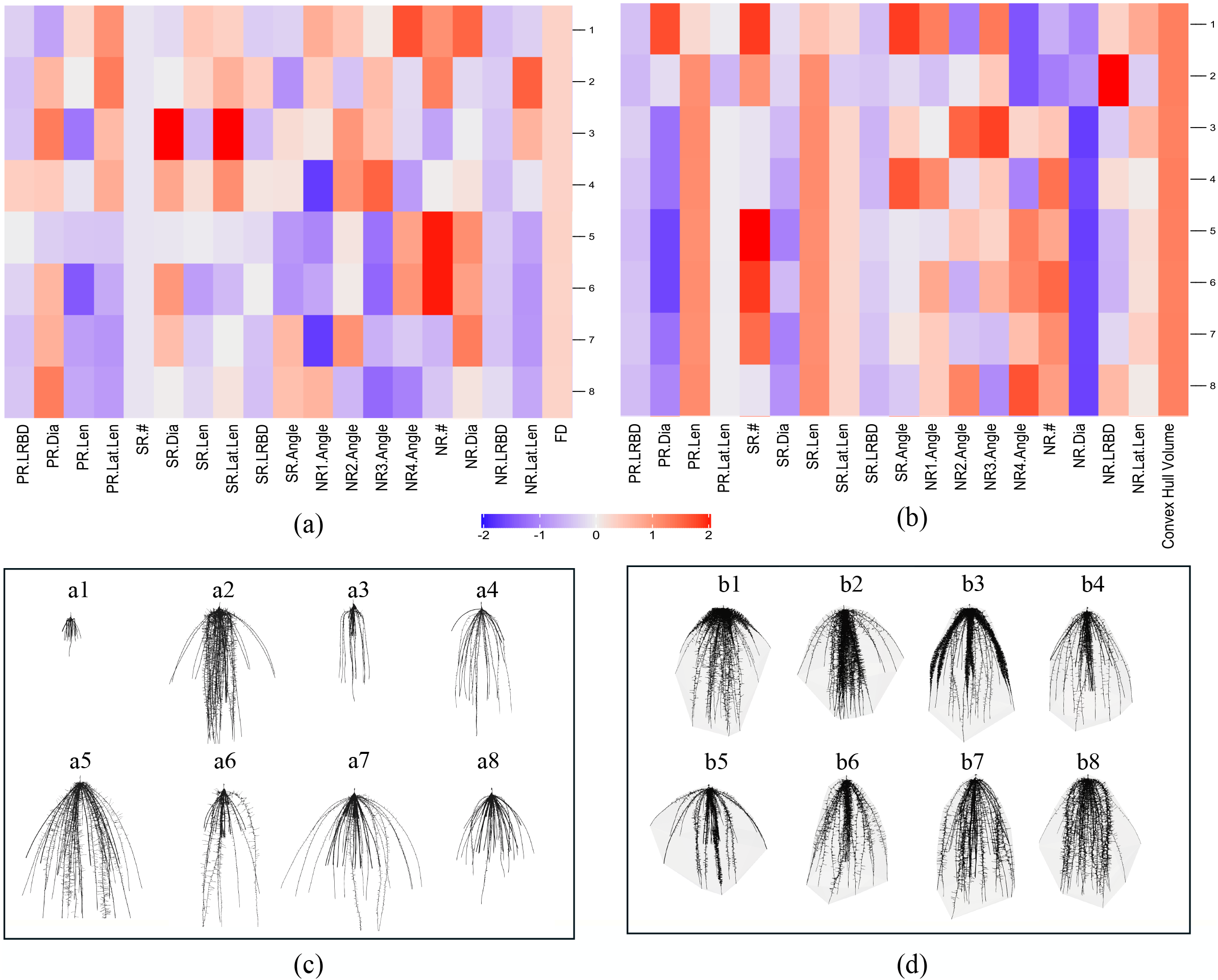
Phene values of maize root phenotypes with comparable FD (a) and convex hull volume (b). The heatmap shows values of the traits obtained by dividing the values with maximum values of respective traits. Phenotypes with similar FD and similar convex hull volume are visualized in (c) and (d) respectively Phenotypes a1 - a8 have similar FD; Phenotypes b1-b8 have similar convexhull volume; PR-Primary Root; SR - Seminal root; NR - Nodal root; LRBD-lateral root branching density; Len - axial root length; Lat.Len - lateral root length; # - number of axial roots; Dia - diameter; FD - Fractal Dimension. (2) one whorl and deep angle (3) two whorls and fanned angles (4) four whorls and fanned angles. The corresponding phenotypes are visualized in lower panel. FD-Fractal dimension.

**Figure 6:**
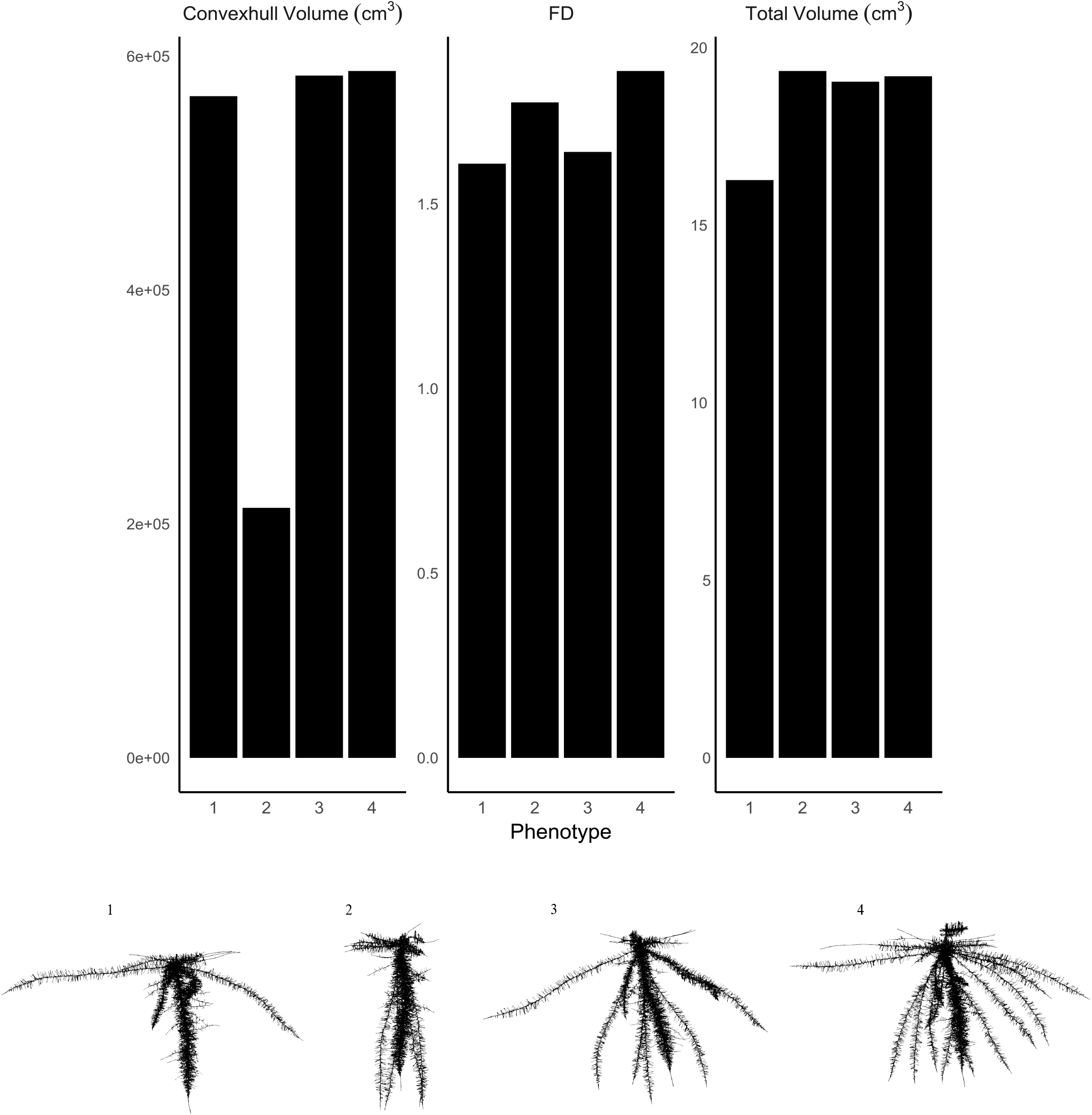
Convex hull volume, FD and total volume of bean root phenotypes with (a) one whorl and shallow angle, (b) one whorl and deep angle(c) two whorls and fanned angle (d) four whorls and fanned angles. The corresponding phenotypes are visualized in lower panel. FD – fractal dimension

### 3.5 Variation in estimates of phene and phene aggregate metrics obtained from 2D projections

In order to study which metrics are not accurately represented by 2D projections, elementary and aggregate phenotypic metrics were estimated from 2D projections obtained by rotating the root system through 360 degrees at 20 degree intervals. It should be noted that *convex hull volume* and area of a 2D projection corresponds to surface area of a 2D hull and the length of the perimeter of a 2D hull respectively. Analysis with 2D image series shows that among phenes, estimates of root growth angle differ when projections are obtained at different rotations. Among aggregate phenotypic trait metrics, the metrics which have angle as one of the most important variables, including *convex hull volume, convex hull area, minor ellipse axis, major ellipse axis, ellipse aspect ratio, solidity, FD* and *FA*, as determined in the random forest analysis, are sensitive to projection. These phenotypic metrics had a coefficient of variation of 10-20 % but some had much greater CV depending on the phenotype in both the maize and bean (Figure 7(a) and Figure 7(b)). The differences in estimates inflated when an aggregate phenotypic trait was calculated as a function of two metrics which are already subject to lot of measurement variation (Figure 7(a) and Figure 7(b)).

**Figure 7:**
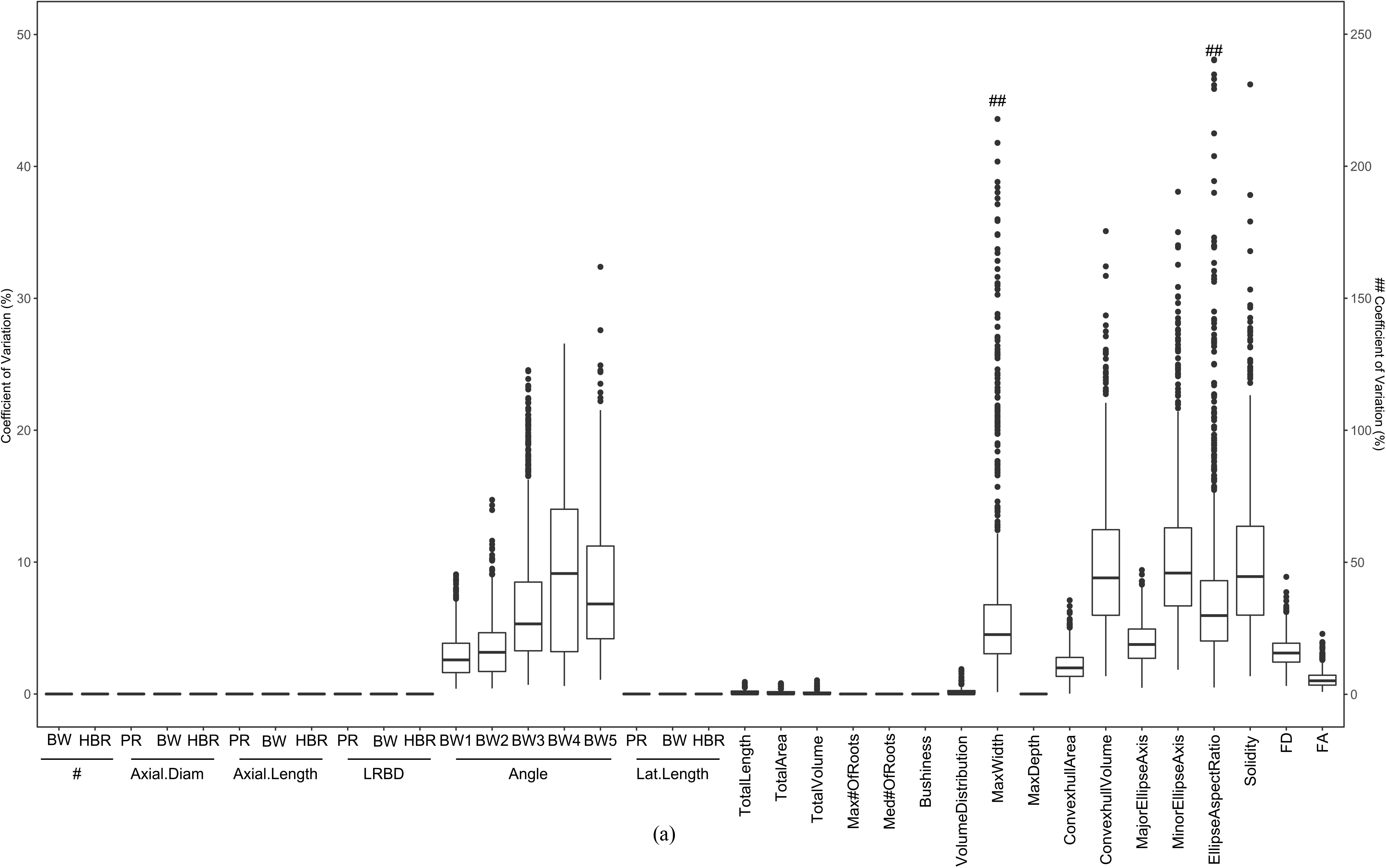
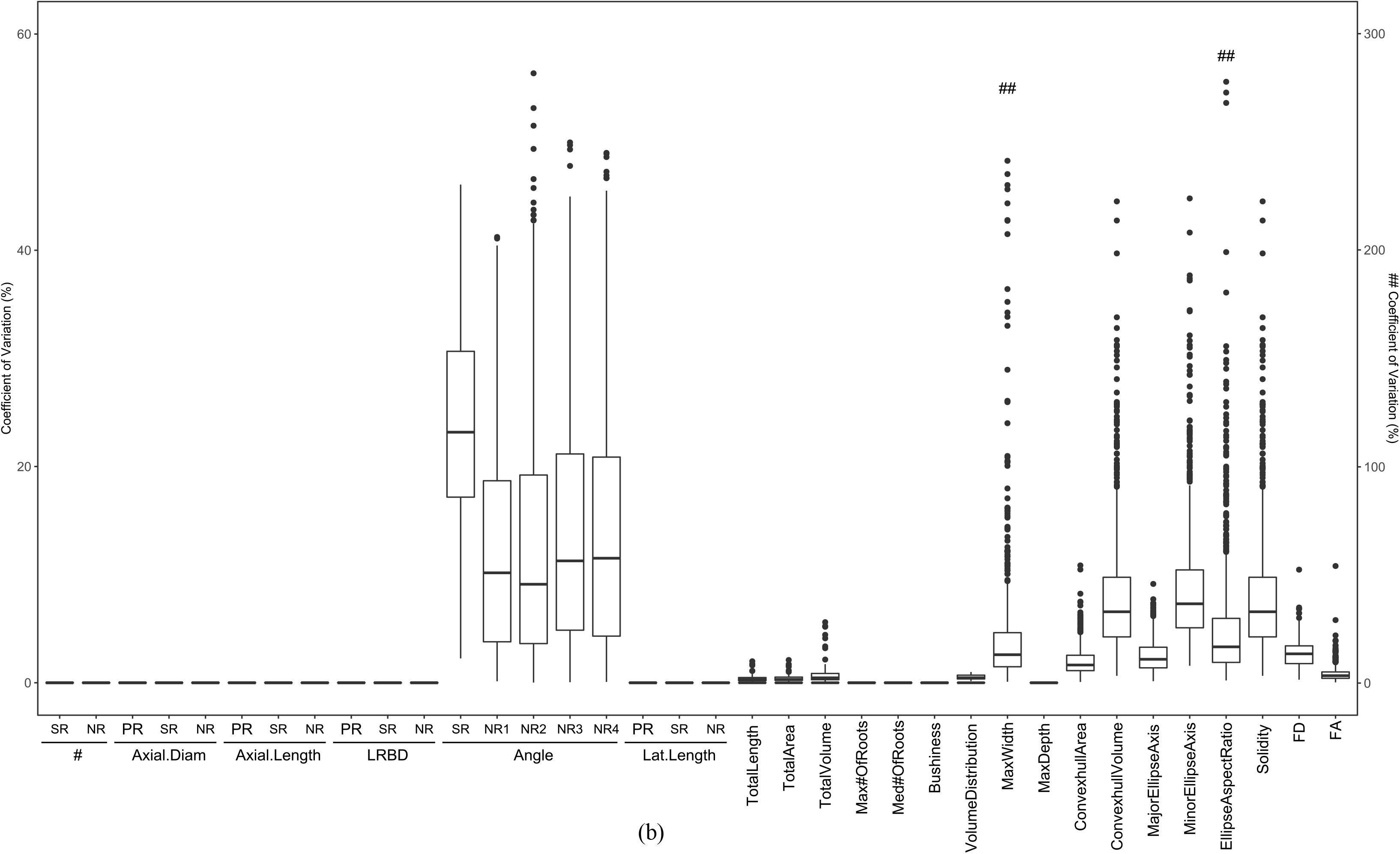
Variation in phene and phene aggregate metrics estimated from rotational series of 2D projected images of 3D root systems of bean (a) and maize (b). BW-Basal root; HBR -hypocotyl-borne root; PR - Primary root; SR-Seminal root; NR Nodal root; # - number of axial roots; Axial.Diam - axial root diameter; Axial.Length - axial root diameter; LRBD - lateral root branching density; Lat.Length - lateral root length; FD - Fractal Dimension; FA Fractal Abundance.

### 3.6 Variation in estimates of phene and phene aggregate metrics over time

Some phene aggregates such increase substantially over 30 days, while some remained relatively static and estimates of some aggregate metric decreased with time (Figure 8(b), Figure 9(b)). Of the traits, *total length, total area, total volume, maximum depth, convex hull area, convex hull volume, major ellipse axis, minor ellipse axis* and *FA pro*gressively increased over time in both bean and maize (Figure 8(b), Figure 9(b), Supplementary Figure S4(b), Supplementary Figure S5(b)). There was only a small change in the *maximum number of roots* in bean over time but this value increased significantly in maize over time (Supplementary Figure S5(b)). The pattern of changes in *FD* over time was dependent on the phenotype. There was a small decrease in *bushiness index* of bean over time (Figure 8(b)). In maize, the phenotypes showed a significant increase from day 10 to 20 followed by a drop from day 20 onwards (Figure 9(b)). The magnitude of increase was dependent on the phenotype. *Volume distribution* was either static or there was a slight increase in the bean phenotypes over time (Supplementary Figure S4(b), Supplementary Figure S5(b)). In maize the change in magnitude of *volume distribution* over time was dependent on the phenotype.

**Figure 8:**
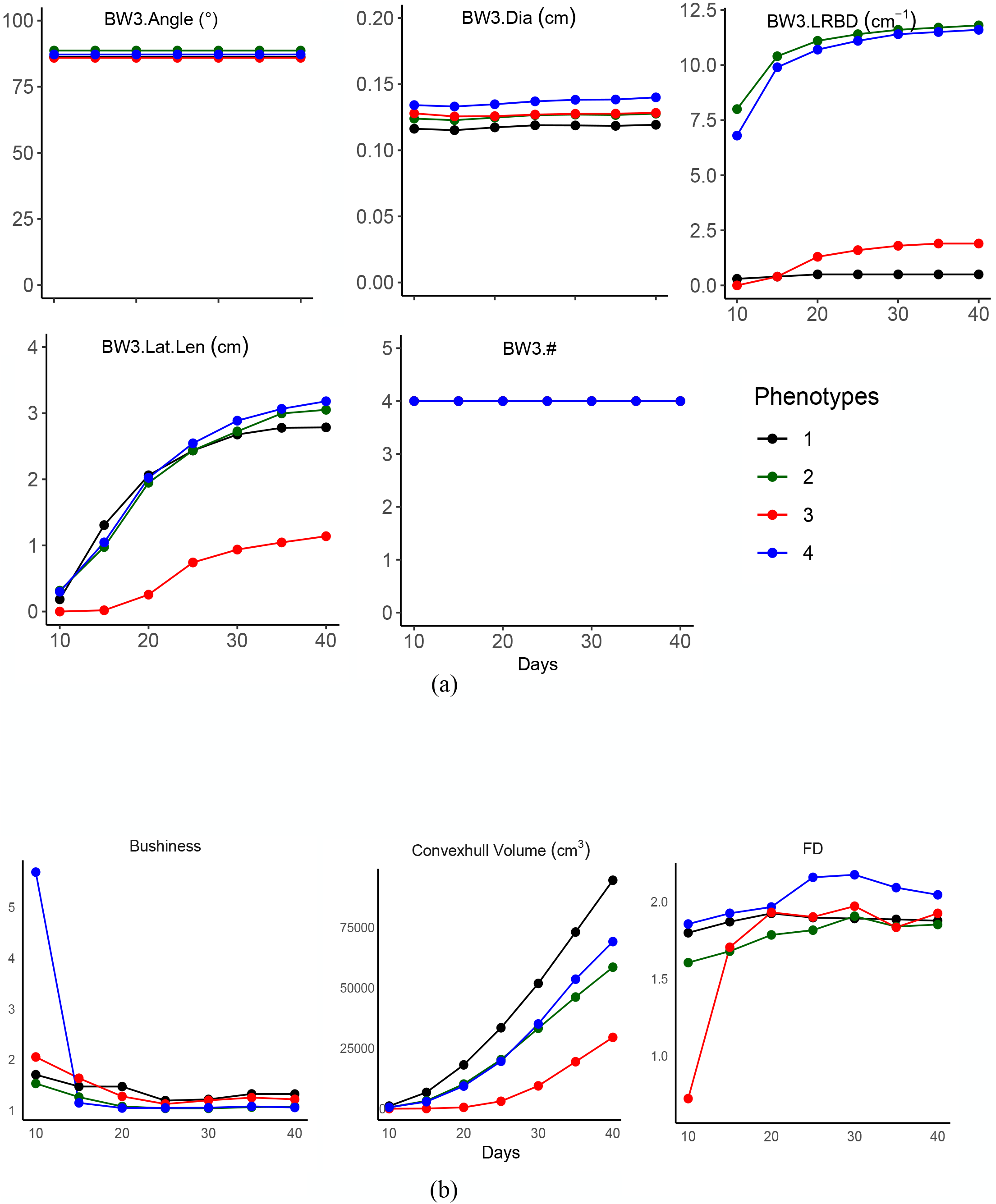
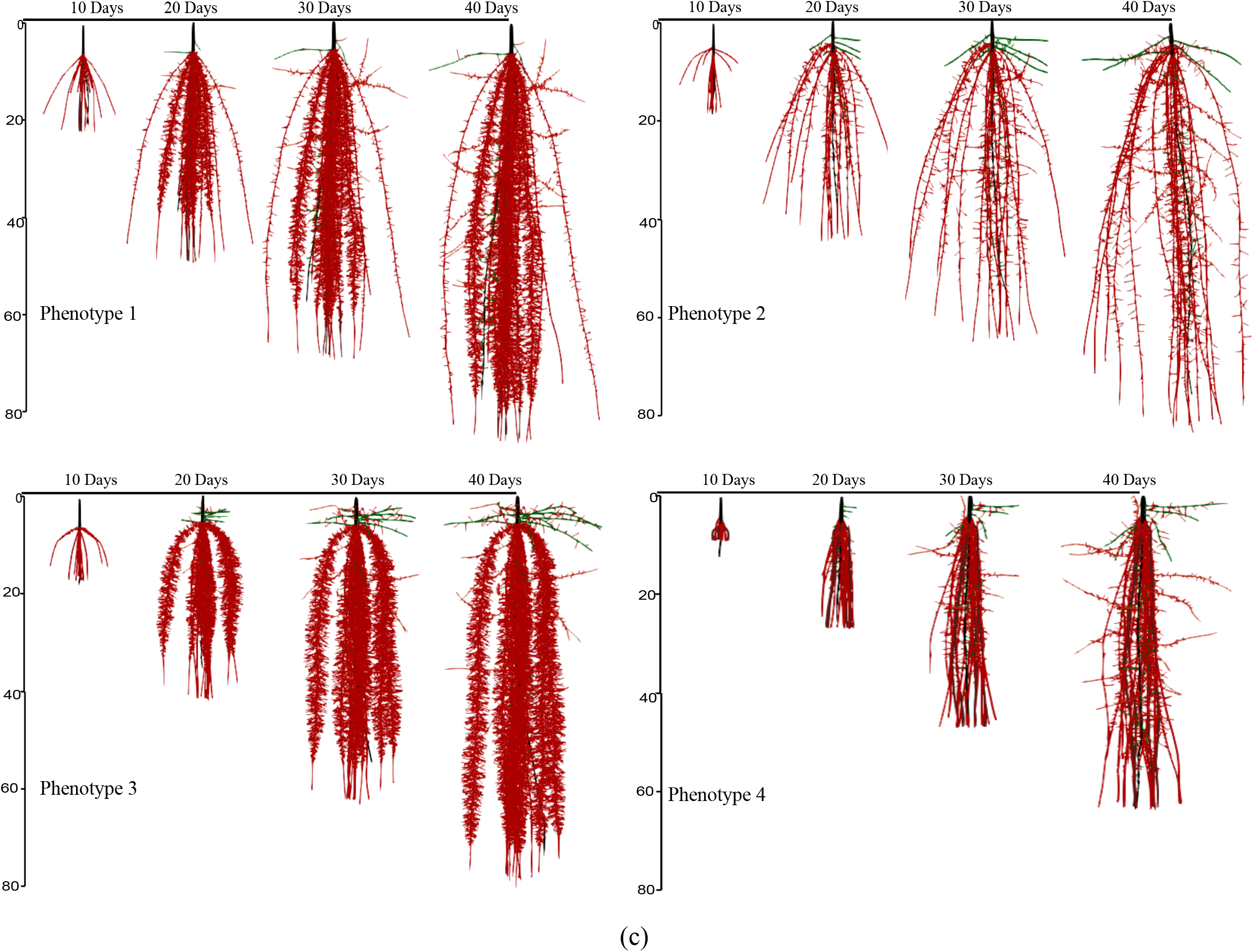
Trait dynamics of bean root phenotypes over 30 days of growth from day 10 to day 40. Change in estimates of phenes associated with basal whorl 3 (BW3) are shown in Figure 8(a). Similar trends were seen in other root classes (Supplementary Figure 4(a)). Change in estimates of the phene aggregates bushiness index, convexhull volume and fractal dimension (FD) are shown in Figure 8(b). Trends in the estimates of other phene aggregates included in this study are shown in Supplementary Figure 4(b). The phenotypes for which the metrics are presented in Figure 8(a) and Figure 8(b) are visualized in Figure 8(c). Primary roots are in black; basal roots in red; hypocotyl-borne roots in green. BW3 - basal roots at whorl 3; Dia - axial root diameter; LRBD - lateral root branching density; Lat.Len - lateral root length; # - number of axial roots.

**Figure 9:**
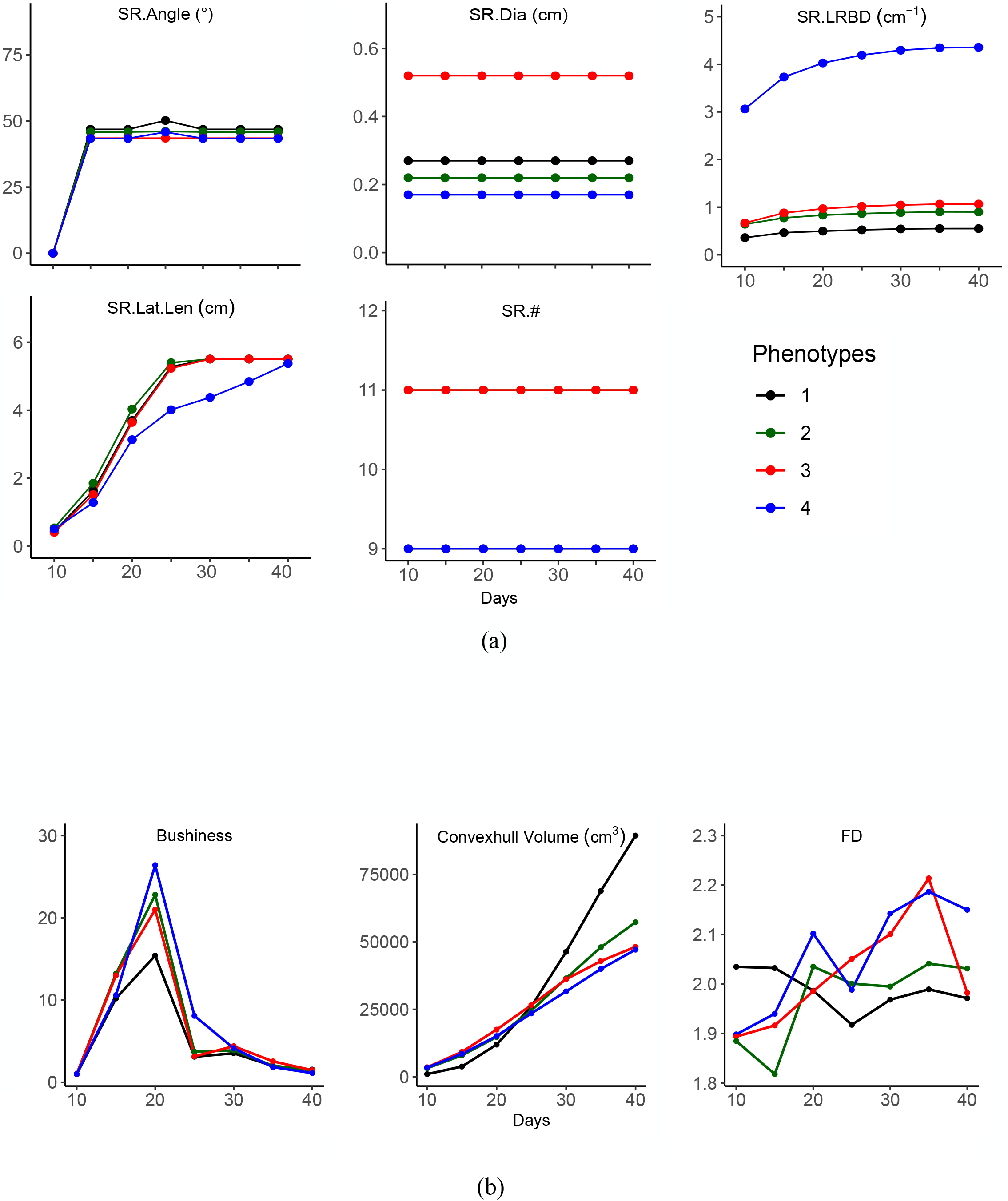
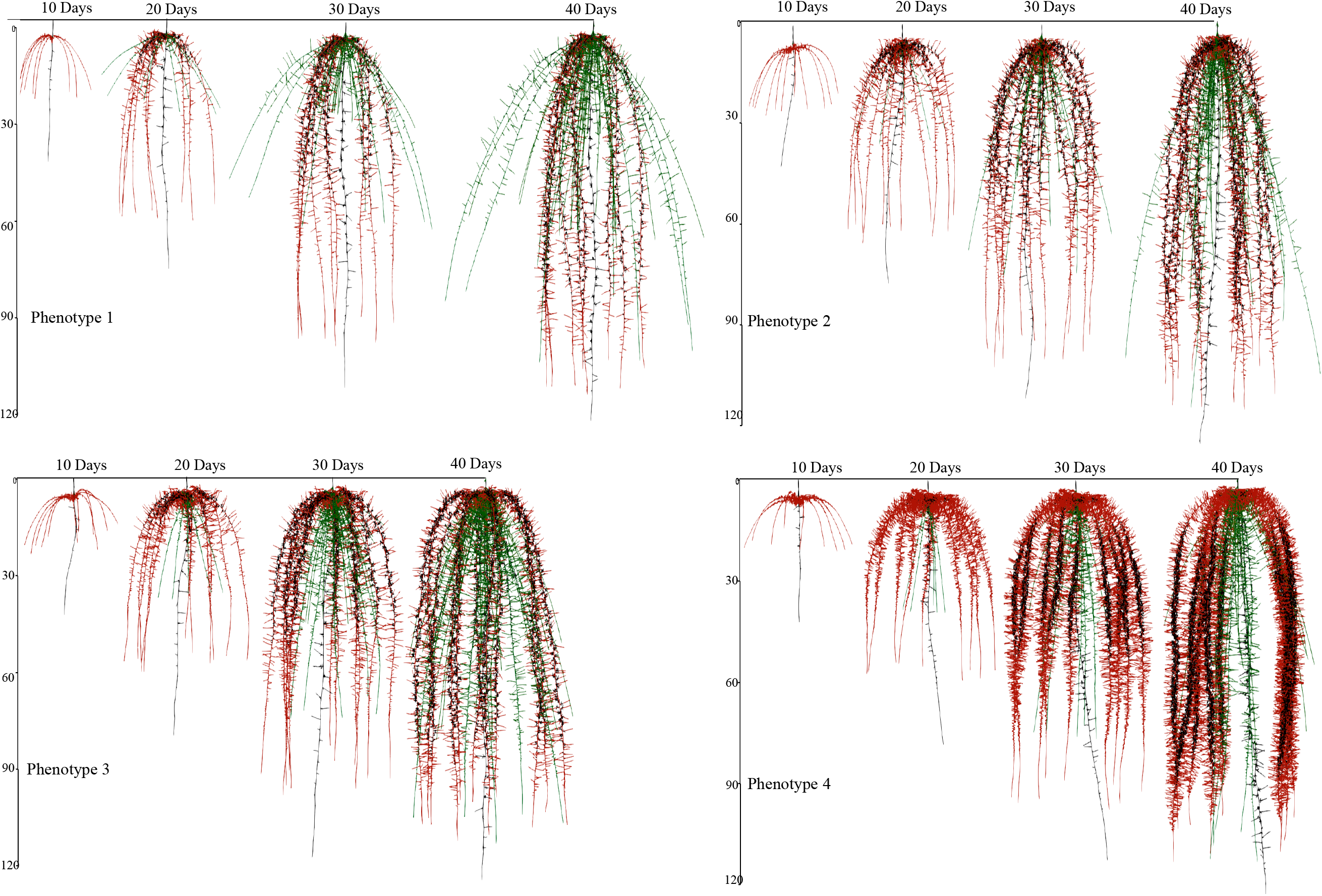
Trait dynamics of maize root phenotypes over 30 days of growth from day 10 to day 40. Change in estimates of phenes associated with seminal roots (SR) are shown in Figure 9(a). Similar trends were seen in other root classes (Supplementary Figure 5(a)). Change in estimates of the phene aggregates bushiness index, convexhull volume and fractal dimension (FD) are shown in Figure 9(b). Trends in the estimates of other phene aggregates included in this study are shown in Supplementary Figure 5(b). The phenotypes for which the metrics are presented in Figure 9(a) and (b) are visualized in Figure 9(c). Primary roots are in black; seminal roots in red; nodal roots in green. SR - seminal roots; Dia - axial root diameter; LRBD - lateral root branching density; Lat.Len - lateral root length; # - number of axial roots.

## Discussion

This study investigated the importance and utility of phenes and phene aggregate traits in phenotyping studies. Our results confirm that phenes are robust and stable over time and also sensitive enough to discriminate between highly similar root systems. In contrast, since phene aggregates capture combinations of subtending phenes, and several combinations of phenes in different states can produce phenotypes which have comparable estimates of phene aggregates, the estimates of phene aggregates are not unique representations of the state of the underlying phenes. Aggregate phene metrics are not stable over time, mostly because there is a rapid development of many elementary root phenes over time. When the number of phenes estimated by the aggregate metric increases, the complex interactions among phenes result in the same phenotype having vastly different estimates for the same aggregate metric at different time points.

### 4.1 Root models can aid exploration of root phenomics

In this study we use *SimRoot* to simulate root systems and use the simulated phenotypes to evaluate various root phenotyping metrics. We used modelling for this study due to constraints in obtaining empirical data caused by limitations in phenotyping methodologies and artifacts due to technicalities in image processing. Phenotyping efforts represent a compromise between throughput, precision and data processing. Many high-throughput phenotyping methodologies involve obtaining 2D metrics and depend on growing plants in controlled growth systems such as pouch, pots, gel plate systems, germination paper, etc. where root architecture is affected due to spatial growth constraints, in particular, branching angles. Not all 3D RSA estimates can be obtained by series of 2D image data; some phenotyping metrics such as volume of non-convex shapes cannot be obtained from 2D projections, especially from complex root systems. Occlusions in 2D images caused by crossing roots increase complexity of systems and reduce accuracy of many 2D estimates; this is especially true for mature root systems which are complex branched structures composed of overlapping and crossing segments (Lobet et al., 2017) ; 3D estimates are better for measuring these “traits” but are biased for other parameters such as surface area due to technicalities in image reconstructions. 3D imaging techniques such as x-ray computed tomography (µCT) and magnetic resonance imaging allow non-invasive studying of spatiotemporal dynamics of root growth (Mooney et al., 2012; Tracy et al., 2012; Schulz et al., 2013; Metzner et al., 2015), but require elaborate data processing and are suitable for relatively small and young root systems due to technical restrictions in container size (Bucksch et al., 2014; Landl et al., 2018) and are scanned at low throughput (Downie et al., 2015; Landl et al., 2018). Studies under controlled conditions enable study of growth of roots over time, however are generally used to assess less complex root structures on younger plants from germination to ca. 10 day after germination (Clark et al., 2011). This is a particular limitation for monocot roots which develop more axial roots over time. Destructive field sampling methods such as shovelomics (Trachsel et al., 2011; Burridge et al., 2016) allow the measurement of the root crown phenotype however is associated with loss and possible displacement of fine roots (Pagès and Pellerin, 1994; Pellerin and Pagès, 1994). Estimates of phenotyping metrics such as fractal dimension is sensitive to incompleteness of the excavated root network (Nielsen et al., 1999; Bucksch et al., 2014).

*SimRoot*, a functional-structural plant model has been used extensively for elucidating the functional value of one or more phenes, and to analyze phene interactions and root complexity (Walk et al., 2004; Walk et al., 2006; Lynch, 2007; Postma and Lynch, 2011a; Postma and Lynch, 2011b; Postma et al., 2014; Dathe et al., 2016; Rangarajan et al., 2018). Simulations with *SimRoot* enable comparing genotypes that vary only in the phene of interest, i.e. near-isophenic lines, which are exceedingly difficult to obtain empirically (Lynch, 2011; York et al., 2013; Rangarajan et al., 2018). A significant advantage of using *SimRoot* is that root architecture over time is known in its entirety devoid of measurement and sampling error. Highly complex root systems can be simulated and resulting root images can be used without any requirement of cleaning images as there is no image noise. Root image co-ordinates are recorded as they grow in 3D space, and so root phenotyping traits can be measured at any time step for any number of time steps without additional effort. One of the major hurdles in phenotyping roots is that artifacts may be present so that the representation of the root system may not be accurate.

### 4.2 Correlation among estimates of phenes and phene aggregates are an emergent property of SimRoot

Our studies with phenes and phene aggregates show that some phenes are highly correlated with each other. *SimRoot* is a mechanistic model and has no fixed relationships for the root architectural parameters. The phenotype is simulated based on a set of input parameters including number of roots of different root classes, root growth angles, root diameter, lateral root branching density with some stochasticity included in each of the parameters. Due to carbon feedbacks and restricted carbon availability, not all phenotypes are simulated. The root system develops based on carbon availability as determined by availability in the seed initially. Plant growth and development occurs as emerging from underlying processes such as photosynthesis, allocation of assimilates, uptake of nutrients and determine the growth of the plant root system (Walk et al., 2006; Postma et al., 2014; Rangarajan et al., 2018). There are no correlations built into the model and the correlations seen among the phenes in the phenotypes are a result of the mechanistic processes that are captured in the model. For example, larger diameter root axes result in larger carbon sinks leaving few resources for other roots. A set of carbon allocation rules determine carbon allocated to different root classes with axial roots having precedence over lateral roots. This is seen as a reduction in lateral root length when the number of roots is greater or when the root diameter is greater. Growth rates of the root tips are a function of carbon availability and if severe carbon limitations occur (as would occur if the phenotype being simulated had many axial roots, greater branching density or large diameter roots or combination of these), axial root length is affected and in extreme cases may inhibit the emergence of roots emerging later. Attempts to factorially design phenotypes based on discrete values of the phene states resulted in some phenotypes not developing for more than few days due to carbon limitations. This is because *SimRoot* keeps track of resource allocation (C, N, P) and trade-offs in carbon allocation result in trade-offs among root traits, as occurs with real plants. The trade-offs include longer axial roots and longer lateral roots when number of axial roots/axial root diameter is reduced, which are seen as high correlations among those phenes. Only those phenotypes that supported plant growth for 40 days were used so that the metrics were dependent only on the phenotype. All metrics were recalculated/extracted from the simulated root system in order to get an accurate estimate of the phenotypic metric.

Correlations also exist among phene aggregates*; maximum depth* and *major ellipse axis* were highly correlated; *Convex hull area, convex hull volume, maximum width* and *minor ellipse axis* were also highly correlated as seen in several other studies *Major ellipse axis* and *maximum depth* are measures of rooting depth (Wedger et al., 2019) and were correlated with primary root length. *Maximum width, minor ellipse axis* and *convex hull* are phene aggregates which characterize expansion in sense of the outer shape of the root system (Paulus et al., 2014). *Maximum width* and *minor ellipse axis* estimates are one-dimensional metrics, *convex hull* is a function of all three dimensions (Mairhofer et al., 2013). These differences mean that as the root grows, estimates of the *convex hull* have a much greater increase in magnitude than does *maximum width*. *Solidity*, which is a ratio of the *total volume and convex hull,* could increase or decrease as *total volume* is dependent on number of roots, lengths of the roots of different root classes and diameters, however *convex hull* estimates the volumetric expansion of the outer shape of the root system.

### 4.3 Phene aggregate metrics are not an unique estimate of phenotype

Phene aggregate measures such as rooting depth are functionally useful traits, as has been demonstrated by several studies. Rooting depth however is influenced by several phenes including root angle, number of roots, LRBD, as shown by several studies (e.g. Manschadi et al., 2010; Trachsel et al., 2013; Saengwilai et al., 2014b; Zhan et al., 2015; Gao and Lynch, 2016). A measure of rooting depth however does not provide any information on the constituent phenes such as rooting angle, number of roots etc. which all contribute to rooting depth. The same is true for other phene aggregate measures such as convex hull volume. Convex hull, defined as the shape of an object created by joining its outermost points, has been used as an indicator of the extent of soil exploration. Calculating convex hull from point clouds requires minimal preprocessing, making it a popularly used phenotyping metric. Although convex hull can provide interesting information about the overall root system shape (Ingram et al., 2012; Zurek et al., 2015), it was not found to be useful in discriminating between phenotypes of different populations (Iyer-Pascuzzi et al., 2010). In a study comparing roots in compacted and uncompacted soil where root geometry is severely affected by soil characteristics, convex hull volume differed by a factor of 3 (Tracy et al., 2012). Here we demonstrate that phenotypes with convex hull estimates within as low as 5% of each other can have phenes expressed in distinctly different states.

While the estimate of a single phene aggregate metric might not be useful in discriminating between phenotypes, using multiple phene aggregate metrics can probably be useful. Each phene aggregate trait gives an estimate of the phenotype by capturing different combinations of phenes. *Total length, area* and *volume* give an estimate of the size of the root system by indirectly measuring the number of roots, length of roots and the diameter of the roots. *Convex hull, minor ellipse axis, major ellipse axis, ellipse aspect ratio, maximum width* and *maximum depth* provide information of the extent of the shape by providing a measurement root angle and root length. Estimates of these phene aggregates, even though they distinguish features of the root system and complement one another in important ways (Topp et al., 2013), do not provide any information on the phene states that comprise the phenotype. Studies aimed at finding root traits which discriminate between populations / phenotypes have found that no single phene aggregate trait was important (Zurek et al., 2015). Which traits were key as well as the number of informative traits were highly dependent on differences between RSA and the imaging day (Zurek et al., 2015). Complexity of RSA over time reinforce the necessity of assessing a large number of traits to distinguish between different varieties as well as individual varieties at different ages (Iyer-Pascuzzi et al., 2010; Topp et al., 2013; Zurek et al., 2015). Accuracy of the different metrics is strongly linked to the root phenotypes analyzed as well as their size and complexity.

### 4.4 Variation in estimates from 2D projection images arise especially due to phenes that determine the geometry of the root system

Root angle is an important phene for soil resource capture; studies have shown that shallow root angles are important for capture of immobile soil nutrients and deep root angles for mobile soil nutrients as well as water capture(Zhu et al., 2005a; Omori and Mano, 2007; Uga et al., 2011; Dathe et al., 2013; Lynch, 2013; Miguel et al., 2013; Miguel et al., 2015; Dathe et al., 2016; Lynch, 2019) . Differences in root growth angle result in phenotypes with distinct differences due to trade-offs in the capture of mobile and immobile soil resources and resulting trade-offs in phenes leading to large effects in biomass production (Ge et al., 2000; Dathe et al., 2016; Rangarajan et al., 2018). Our results show that estimate of root angle is affected by the 2D projection of the root system. Root angle determines the geometry of the root system and was found to be an important variable in determining variations in *convex hull area, convex hull volume, maximum width* and *minor ellipse axis* (Table 2). Aggregate phene traits capturing the geometry or overall shape of the root cannot be measured accurately using estimates derived from 2D data. The variation in the estimates of root angle when measured using 2D projections affect the estimates of all phene aggregate traits in which they play an important role directly or indirectly; these include secondary phene aggregate traits such as solidity, ellipse aspect ratio as well as root complexity traits *FD* and *FA* (Figure 8 and Figure 9). Variation is greater in phene aggregates which are estimates of some function of more than one aggregate phene. Even though our root phenotypes are simulated, they are based on empirical parameters, and differences in number of roots, angles of each root class etc. were varied and as a result, our root phenotypes were not symmetrical, to replicate actual root system in fields. This is important because most roots found in nature are not symmetrical. We found that greater asymmetry was associated with greater variation in the aggregate phenotypic metrics estimated from 2D projections. Results from studies using 2D images from gel culture, growth pouches, narrow growth containers with a transparent face, etc., should be interpreted with caution.

### 4.5 Variation in phene aggregate metrics with time is species dependent

We analyzed root phenotypes of two species, maize and common bean, representing a monocot and a dicot root architecture. The main difference between bean, which is a dicot root system, and monocot root systems is that new roots (laterals) emerge from already existing roots in dicots, whereas in monocots nodal roots continually emerge over time from shoot nodes near or above the soil surface (Rangarajan et al., 2018). Therefore, the vertical distribution of roots vary between maize and bean, with the bean root system having a relatively equal root distribution whereas maize has more proportion of roots in the topsoil (Postma and Lynch, 2012; Zhang et al., 2014). The number of roots as well as root diameter depends on the nodal position in maize. This is probably the reason for the great temporal variation in metrics such as volume distribution and bushiness index which are related to root size. It has been suggested that metrics accurate for small dicot root systems might fail for large dicot or small monocot root systems (Lobet et al., 2017). Our study confirms that estimates of phene aggregates are not only dependent on phenotype and time but also on the plant species.

### 4.6 Metrics of root complexity

Fractal parameters are different from all the estimated phene aggregates in that they do not provide information on shape of the phenotype, extent of shape or size of the root system, but instead measure the geometric complexity of the root phenotype (Fitter and Stickland, 1992; Nielsen et al., 1997; Nielsen et al., 1999). All the phenes tested were important in determining fractal estimates. Fractal dimension was useful in differentiating between P inefficient and P efficient bean genotypes (Nielsen et al., 1999) as well study of roots fractal parameters with uptake of diffusion limited nutrients and between genotypic variation in wheat, study developmental responses in rice (Manschadi et al., 2008; Wang et al., 2009). It was found, however, that not a single but combinations of multiple fractal measurements provide useful information (Nielsen et al., 1999; Walk et al., 2004). Phenotypes with comparable aggregate phene trait estimates can be a result of different combinations of phenes in distinctly different phene states. This implies that estimates of phene aggregate traits measure the aggregate of multiple phenes (York et al., 2013). Studies have shown that complex phenotypic traits such as root complexity as measured by fractal analysis are determined by a multitude of genes with small effects (Grift et al., 2011). Even though several studies have resulted in identification of QTLs for aggregate phene traits (Topp et al., 2013; Atkinson et al., 2015; Zurek et al., 2015; Kenobi et al., 2017), only two genes directly controlling RSA have been cloned (Uga et al., 2011; Wedger et al., 2019). Estimates of QTL locations or effects *per se* do not give us direct biological information regarding the product or function of each gene and the interactions among genes (Bernardo, 2008). Phenes are unique, meaning, are the product of only one set of genes and processes at a specified scale of resolution (Lynch and Brown, 2012; Lynch, 2019) and so, phene selection is more genetically tractable than selection for traits that aggregate multiple phenes, because axiomatically phenes are under simpler genetic control than any combination of phenes (Lynch, 2019).

### 4.7 Selection of phenotypes based on phenes are useful for breeding

Several phenes have been studied and their functional utility has been established including number of roots (crown roots in maize, basal roots in bean), root growth angle (shallow for phosphorus uptake and deep rooting angle for nitrogen capture), lateral root branching density and length for nitrate uptake(Zhu et al., 2005b; Lynch, 2013; Trachsel et al., 2013; Saengwilai et al., 2014; Miguel et al., 2015; Zhan and Lynch, 2015; Rangarajan et al., 2018; Sun et al., 2018). In the bean root system, basal roots emerge at the seedling stage and seedling root phenotypes have significant relationships with mature root phenotypes in the bean root system. Number of basal roots as well as basal root growth angle is stable over time as proven by the fact that studies selecting for basal root number and angle at different stages of growth from seedling to few weeks old plants (Liao et al., 2001; Vieira and Lynch, 2001; Vieira et al., 2008) have been consistent. Genetic factors explained 52% to 57 % of genetic variation of phenes in bean including basal root whorl number, basal root number, adventitious root number, and 52% of phenotypic variation in taproot length in seedlings (Strock et al., 2019). Crown root and brace root number, angle and LRBD were found to be genotype-specific and did not change across growth stages in maize (Trachsel et al., 2013). Basal diameter remains constant in maize while apical diameter varies; in dicots like bean, diameter increases with age due to secondary root growth (Strock et al., 2018). Root growth/ elongation rates determine the length of the root and are thought to be phenes (York et al., 2013; Strock et al., 2019). However, carbon limitations could result in delay of emergence of axial roots as well as play a role in determining the final number of axial roots. Demotes-Mainard and Pellerin (1992) have observed on maize that the emergence of axial roots was delayed, and the final number of axial roots was reduced, with increasing levels of competition for light between plants. Time of emergence of roots could also be an important phene, especially in maize where roots emerge from different nodes over time. Recent studies have shown that cellular anatomy varies among nodes providing evidence for node-specific traits (Yang et al, 2019). Our approach using elemental phenes to discriminate between architecturally and anatomically distinct phenotypes based on phene states has been used successfully for selection of functionally superior phenotypes for different crop species (Burridge et al., 2017). We suggest that it is best to study the phenotypes at their elementary level of organization, namely phenes in order to get a better understanding of their functional value in terms of the interactions among the phenes and also to identify their genetic features.

## Conclusions

These results demonstrate that phenes including number of roots, diameter of roots, lateral root branching density and root growth angle provide reliable descriptors of root phenotypes. Phenes are also stable over time and independent of time of phenotyping. Estimates of phenes provide a complete description of the resulting phenotype and also enable easier prediction of functional attributes the phenotype could potentially have. Data from our *in-silico* phenotyping environment provides access to complete information concerning root architectural phenotypes without measurement error, sampling limitations, or confounding factors such as phenotypic plasticity or root loss. Even under these conditions, estimates of aggregate phenotypic metrics are less reliable than those of phene states. Even though the estimates of aggregate phenotypic metrics are dependent on the phenotype, the estimates are not unique estimates of underlying states of the constituent phenes. Estimates of phene aggregates also vary in magnitude at different time points of growth, the magnitude of change being dependent on the aggregate phenotype metric used as well as the constituent phenes. Unlike methods used to estimate aggregate phenotypes, estimation of phenes involves simple, straightforward procedures and yield reliable results. We suggest that measurement of phenes provides data that are more robust, reliable and relevant than metrics that estimate the aggregation of multiple subtending phene states. We show this in the context of root architectural phenotypes but propose that these concepts apply to phenomic analysis of any organism.

## Acknowledgements

HR designed and conducted the research, developed the analytical framework, analyzed the results, and wrote the manuscript. JPL conceived and supervised the project and assisted with experimental design, data interpretation and manuscript preparation. This research was supported by the National Institute of Food and Agriculture, U.S Department of Agriculture Hatch project 4732 and the Foundation for Food and Agriculture Research ‘Crops in Silico’ project. The authors declare that there is no conflict of interest regarding the publication of this article. Data generated in this study is freely available from the corresponding author. The executable code of the version of *SimRoot* employed in this study, parameters used to generate these data, and the raw data, are all available at https://figshare.com/s/58c7599752bcb75fbd76.

## Supplementary Materials

### Supplementary 2

Supplementary Table S1: Range of input values for generating bean root phenotypes. PR – primary root; HBR-Hypocotyl-Borne-Root; BW – Basal Whorl; BW1, BW2, BW3, BW4, BW5 refer to the position of the basal whorl counted from basipetal to acropetal position; Dia – axial root diameter; Lat.Dia – lateral root diameter; LRBD – lateral root branching density.

Supplementary Table S2: Range of input values for generating maize root phenotypes. PR - Primary Root; SR -Seminal Root; NR-Nodal Root; NR1, NR2, NR3, NR4 refer to the nodal root position; Dia – axial root diameter; Lat.Dia – lateral root diameter; LRBD – lateral root branching density. *NR at different positions were considered to have similar parameters.

**Supplementary Figure 1(a)-1(i):**
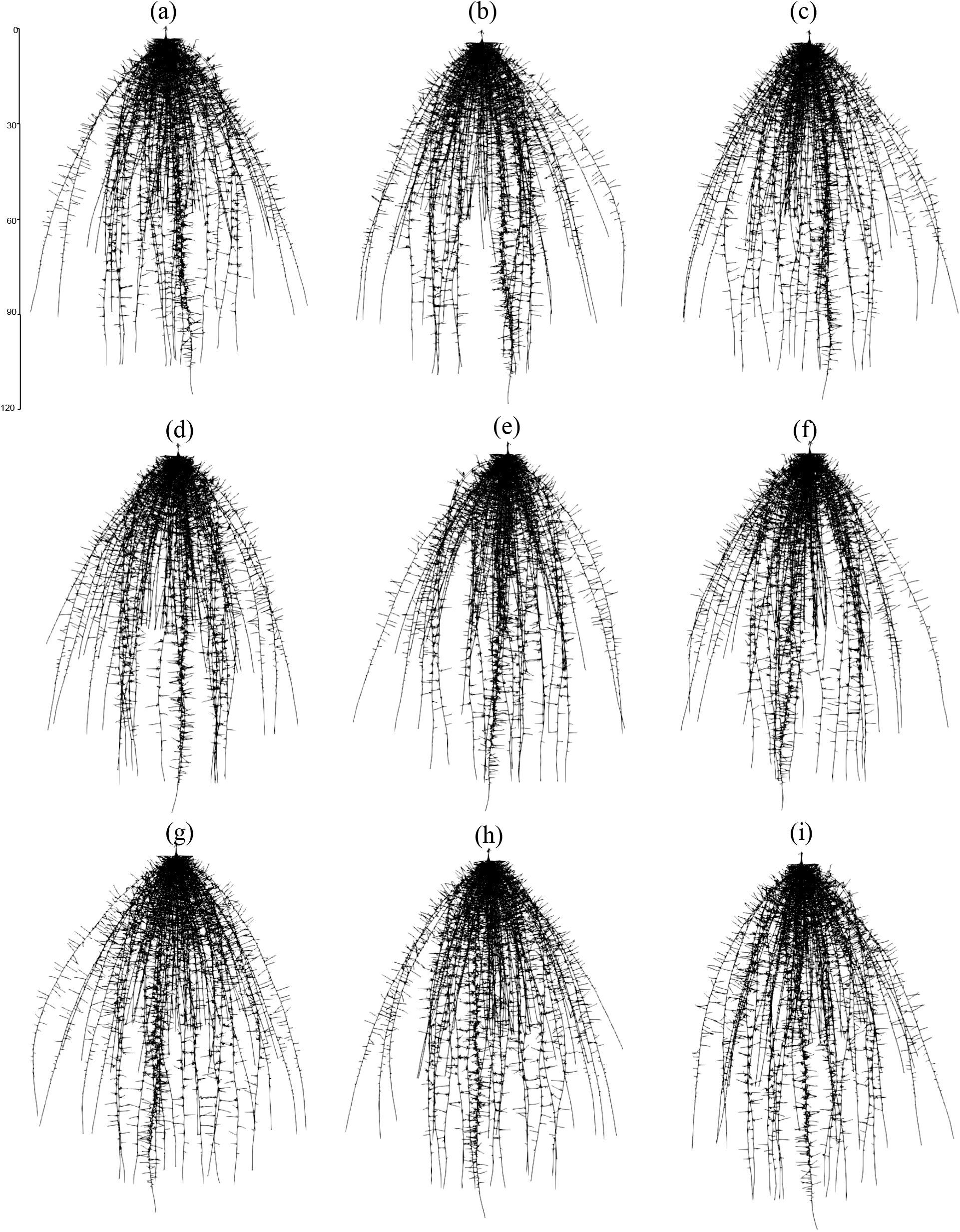
Representative images of 2D projections of a maize root system rotated by 20°, 60°, 100°, 140°, 180°, 220°, 260°, 300°, 340°.

**Supplementary Figure 2:**
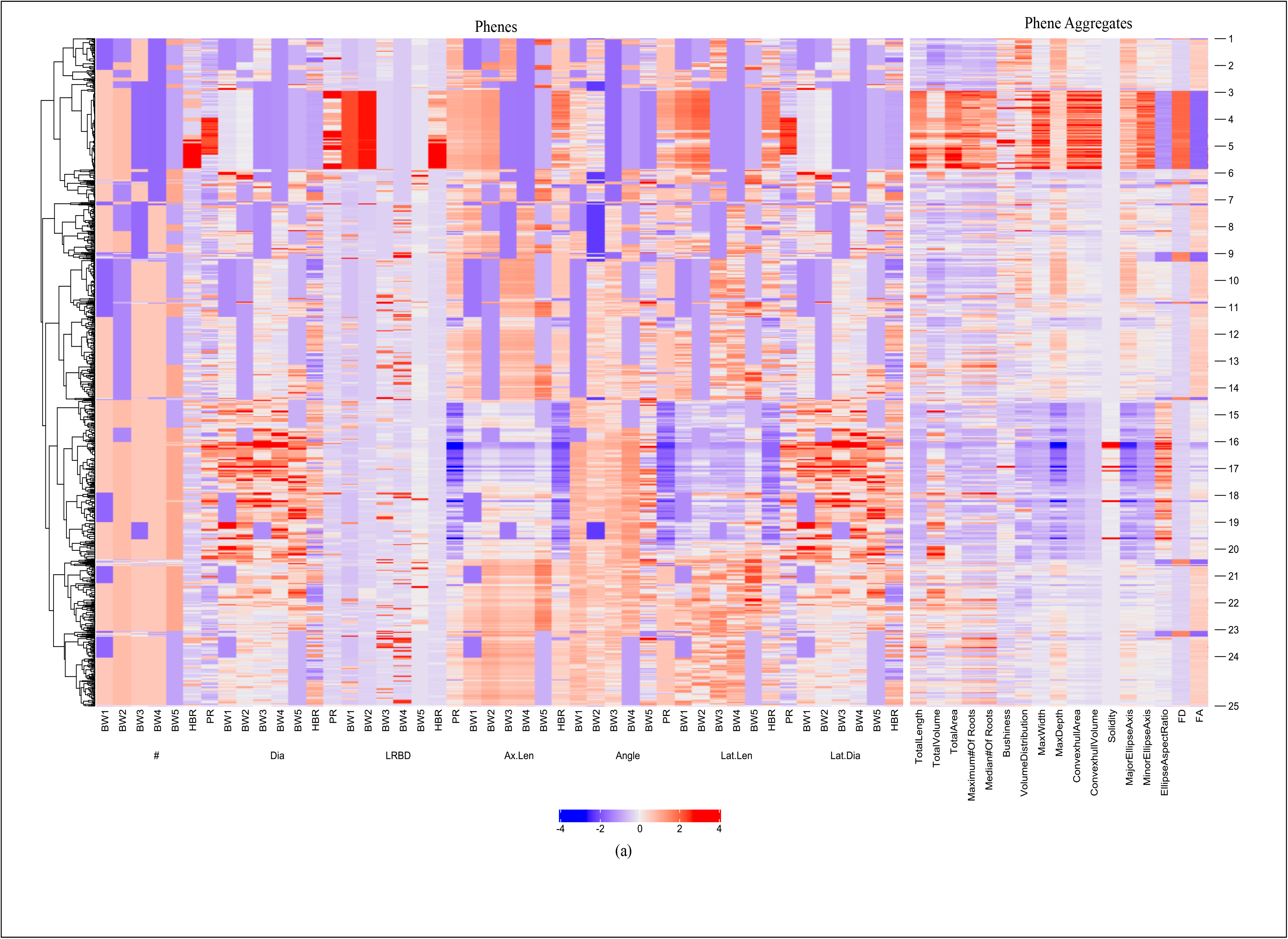
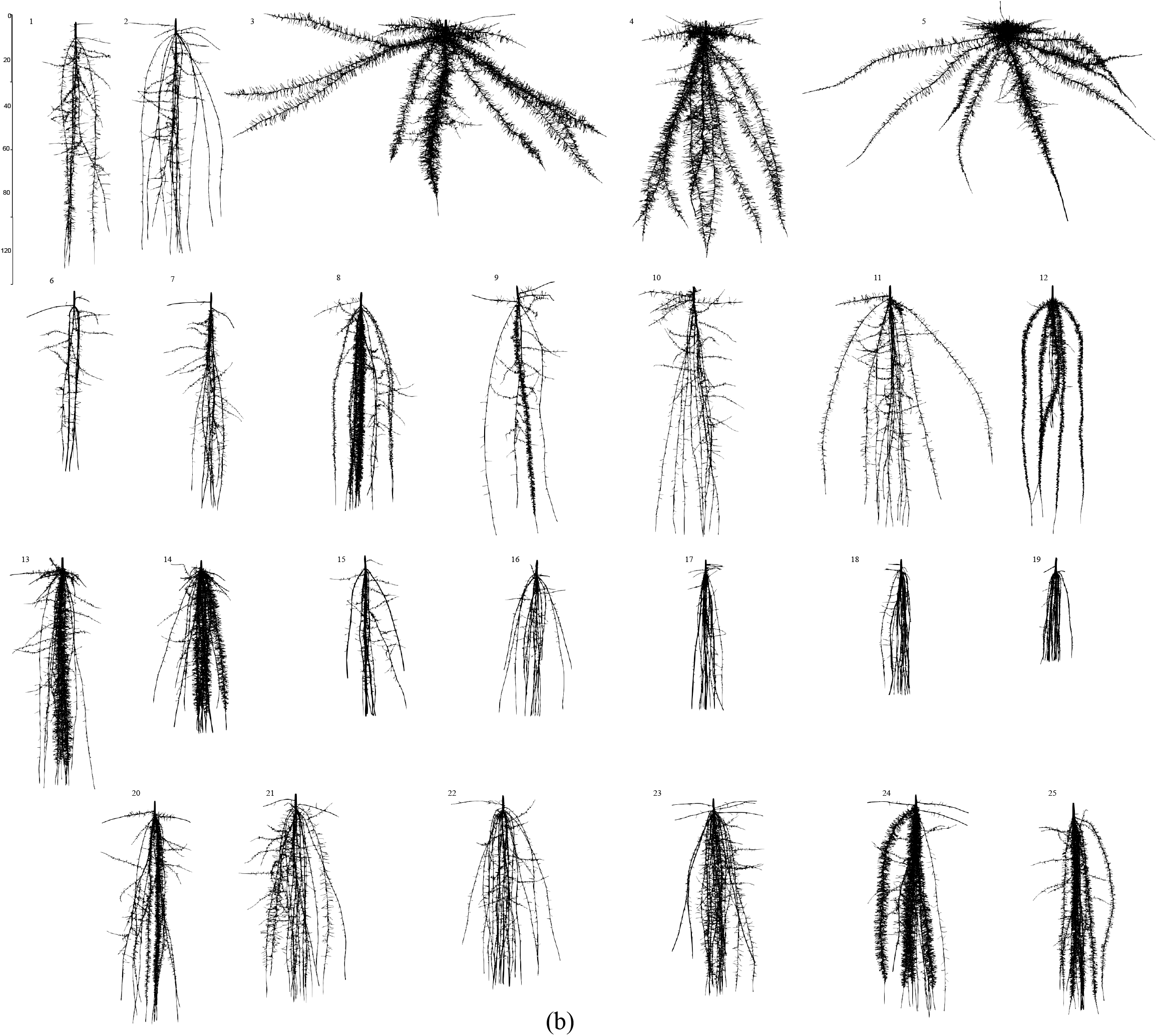
Cluster heatmap of phenotypic traits. Hierarchical clustering of all bean phenotypes was generated using Spearman correlation coefficient of max-min scaled phene values at 40 days (a). The color scale indicates the magnitude of the trait values (blue, low value; red, high value). The numbers indicated on the heatmap refer to a representative phenotype in the specific region of the heatmap. The corresponding phenotypes are visualized in (b). # - Number of roots; Axial.Diam - axial root diameter; LRBD - lateral root branching density; Axial.Length - axial root length; Lat.Length-lateral root length; Lat.Diam - lateral root diameter; BW1 - basal roots at whorl 1; BW2 - basal roots at whorl 2; BW3 - basal roots at whorl 3; BW4 - basal roots at whorl 4; BW5 - basal roots at whorl 5; HBR - hypocotyl-borne roots; PR - primary root.

**Supplementary Figure 3:**
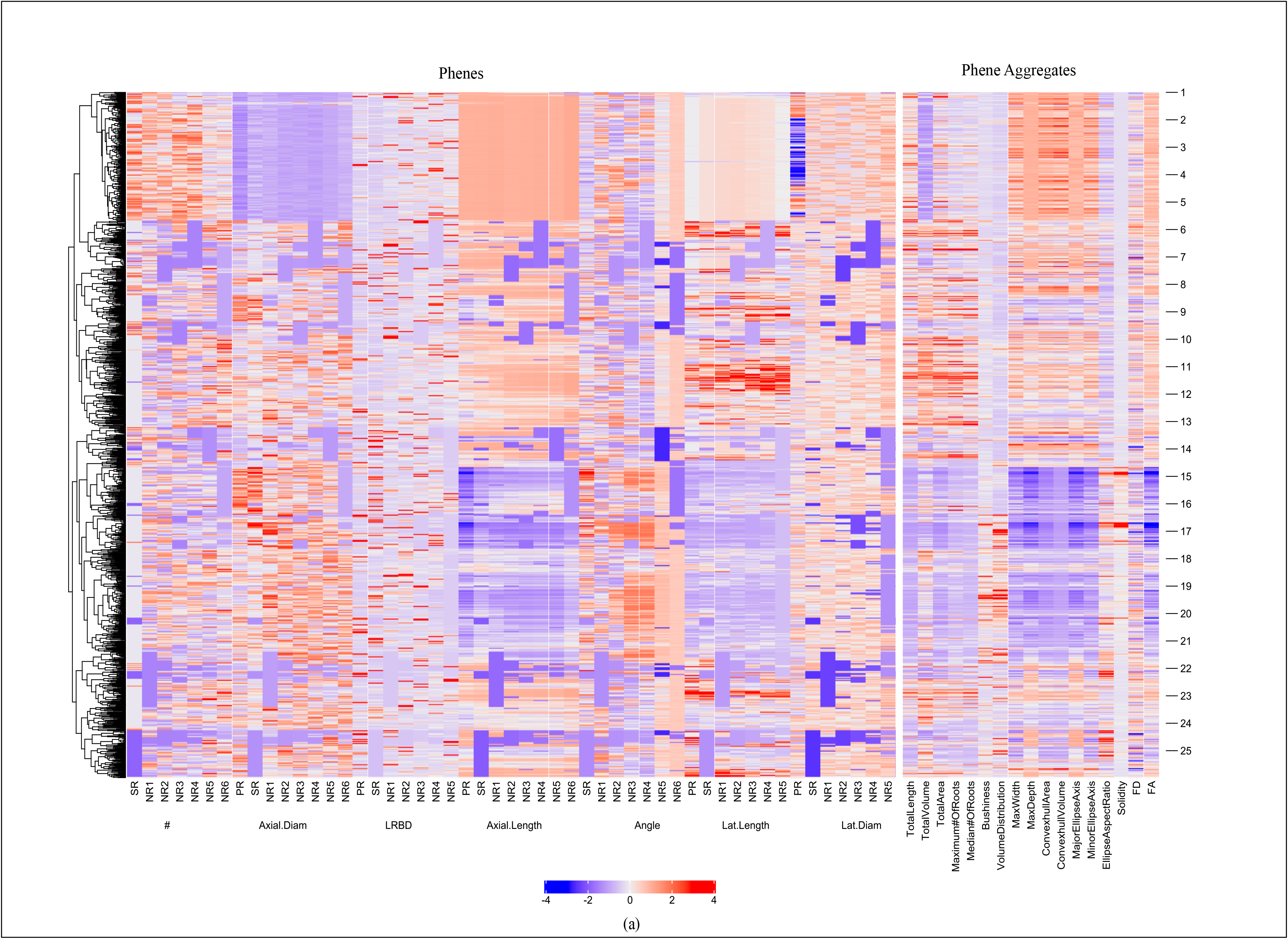
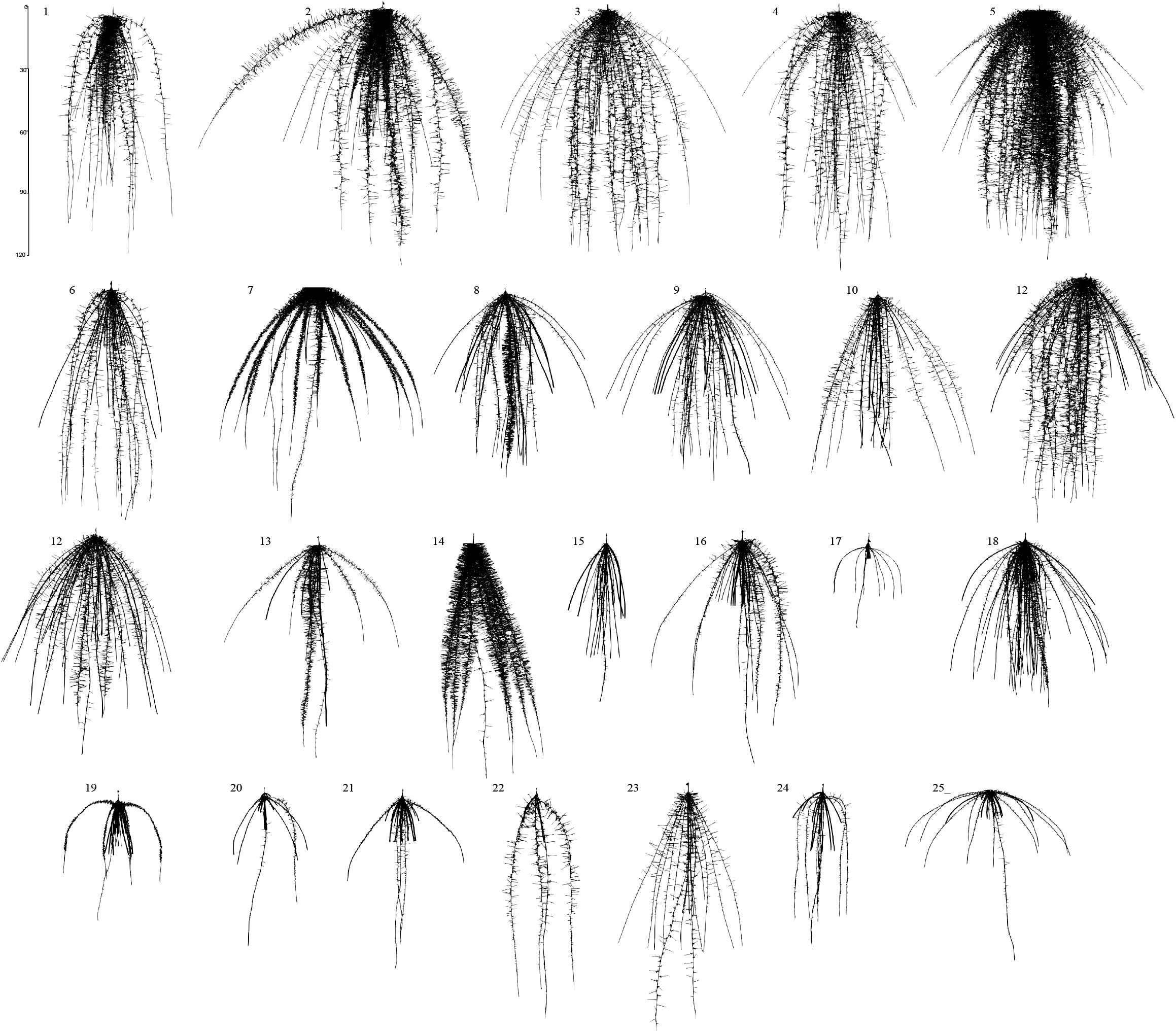
Cluster heatmap of phenotypic traits. Hierarchical clustering of all maize phenotypes was generated using Spearman correlation coefficient of max-min scaled phene values at 40 days (a). The color scale indicates the magnitude of the trait values (blue, low value; red, high value). The numbers indicated on the heatmap refer to a representative phenotype in the specific region of the heatmap. The corresponding phenotypes are visualized in (b). # - Number of roots; Axial.Diam - axial root diameter; LRBD - lateral root branching density; Axial.Length - axial root length; Lat.Length - lateral root length; Lat.Diam - lateral root diameter; NR1 - nodal roots at position 1; NR2 - nodal roots at position 2; NR3 - nodal roots at position 3; NR4 - nodal roots at position 4 ; SR - seminal roots; PR - primary root.

**Supplementary Figure 4:**
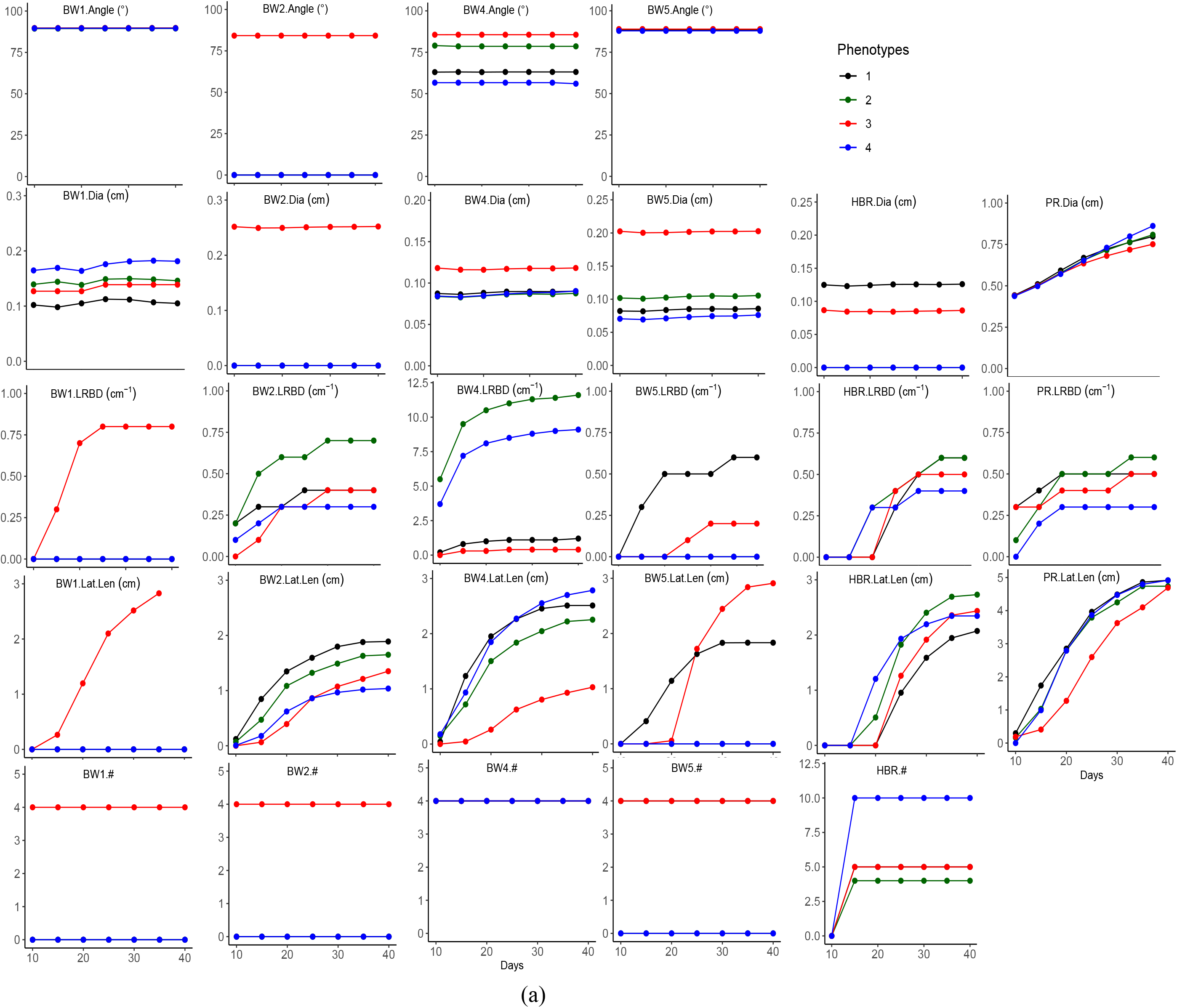
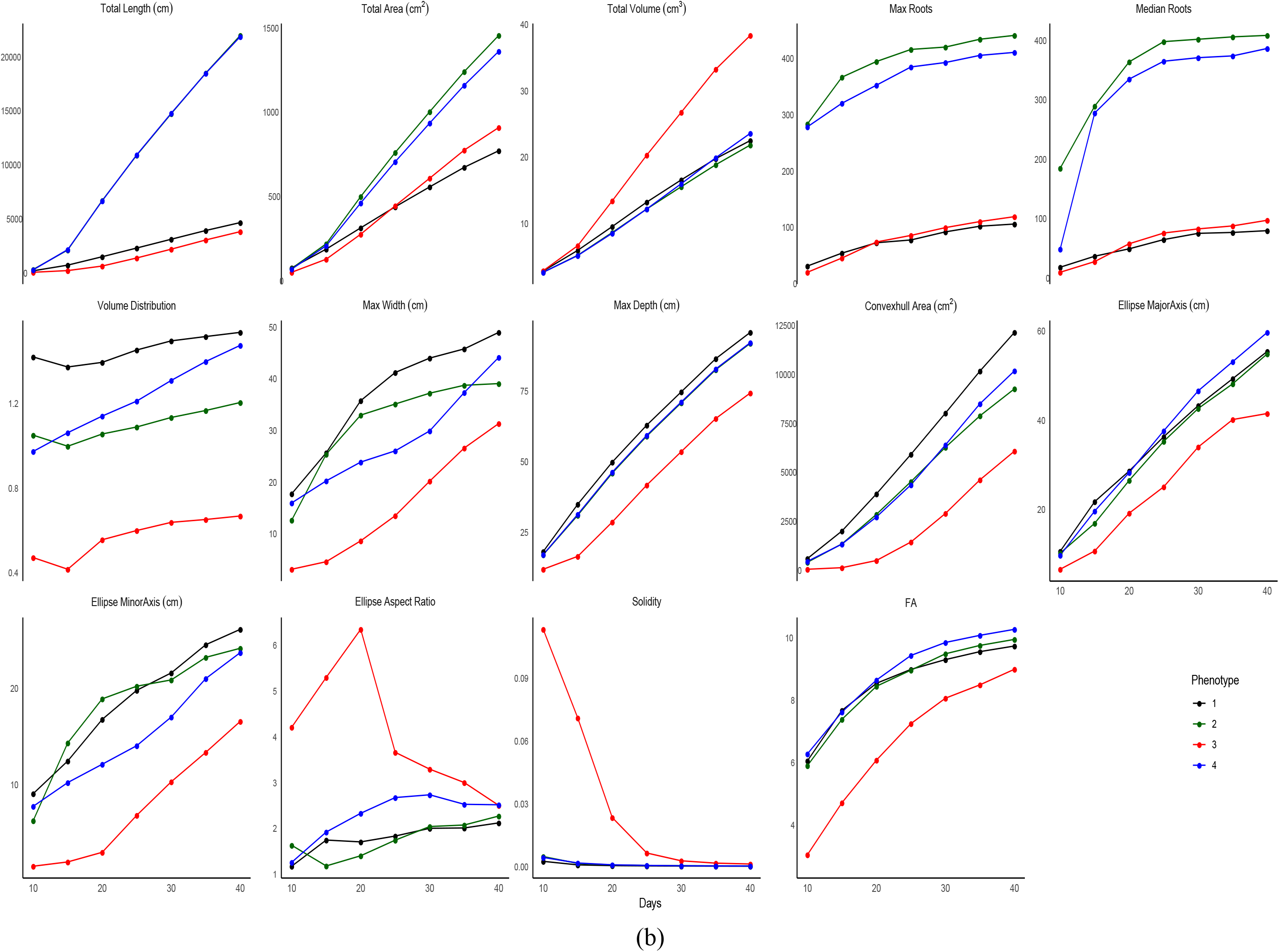
Trait dynamics of bean root phenotypes over 30 days of growth from day 10 to day 40. Change in estimates of phenes (a). Change in estimates of the phene aggregates (b). BW1 - basal roots at whorl 1; BW2 - basal roots at whorl 2; BW4 - basal roots at whorl 4; BW5 - basal roots at whorl 5; HBR - hypocotyl-borne roots; PR - primary root; Dia - axial root diameter; LRBD - lateral root branching density; Lat.Len - lateral root length; # - number of axial roots; FA - fractal abundance.

**Supplementary Figure 5:**
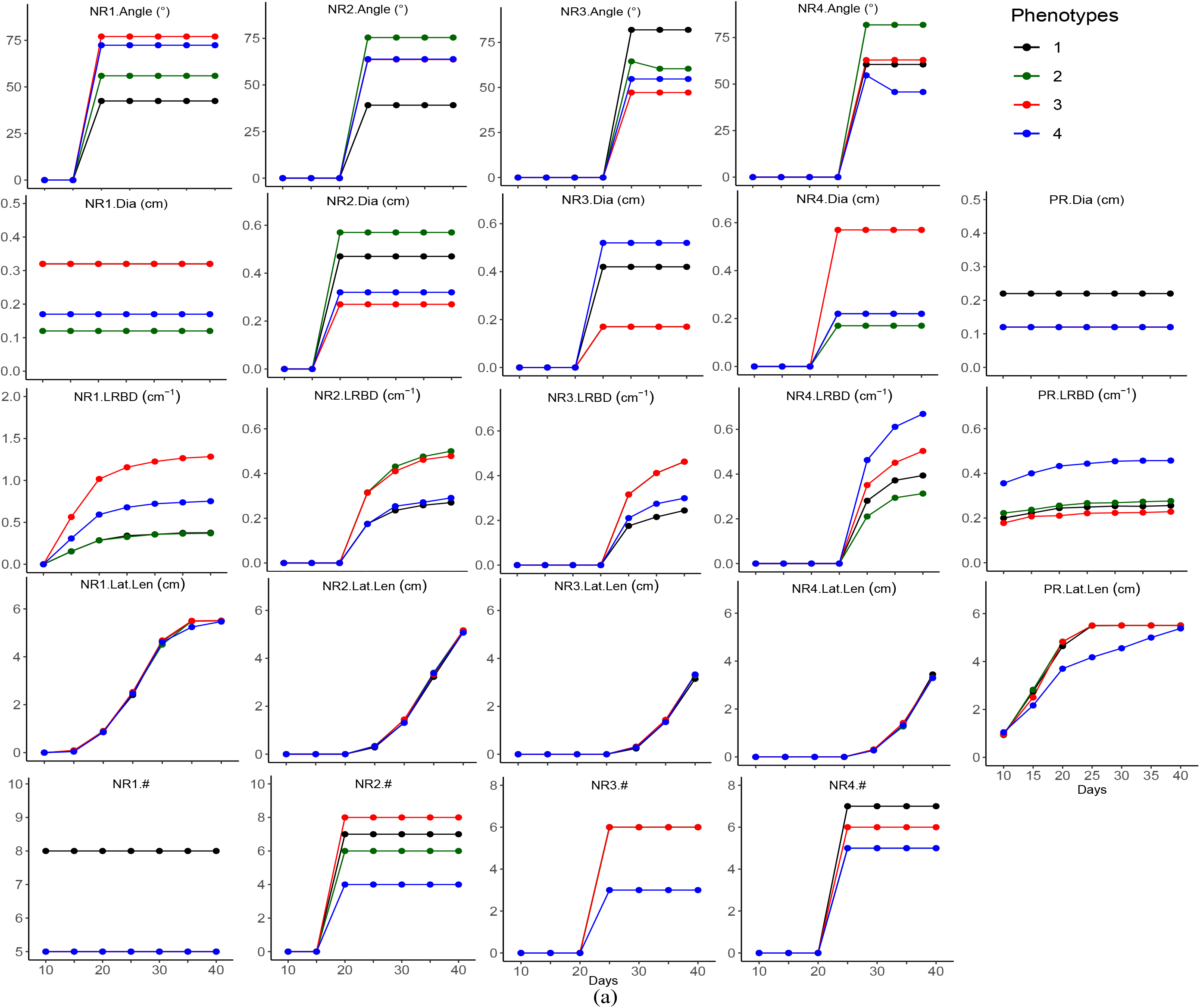
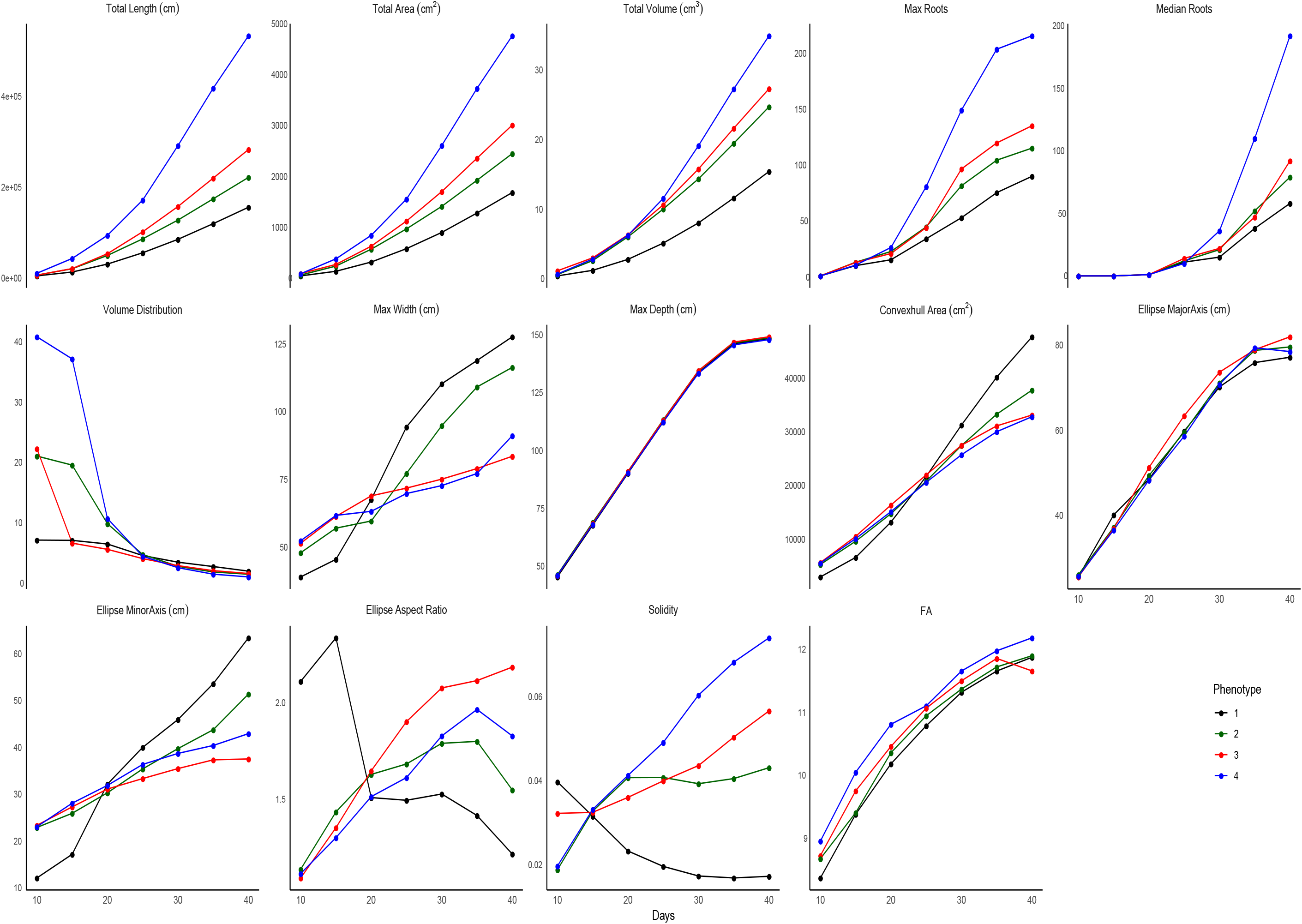
Trait dynamics of maize root phenotypes over 30 days of growth from day 10 to day 40. Change in estimates of phenes (a). Change in estimates of the phene aggregates (b). NR1 - nodal roots at position 1; NR2 - nodal roots at position 2; NR3 - nodal roots at position 3; NR4 - nodal roots at position 4; PR - primary root; Dia - axial root diameter; LRBD - lateral root branching density; Lat.Len - lateral root length; # - number of axial roots; FA - fractal abundance.

